# Decoding neural responses to motion-in-depth using EEG

**DOI:** 10.1101/661991

**Authors:** Marc M. Himmelberg, Federico G. Segala, Ryan T. Maloney, Julie M. Harris, Alex R. Wade

## Abstract

Two stereoscopic cues that underlie the perception of motion-in-depth (MID) are changes in retinal disparity over time (CD) and interocular velocity differences (IOVD). These cues have independent spatiotemporal sensitivity profiles, depend upon different low-level stimulus properties, and are potentially processed along separate cortical pathways. Here, we ask whether these MID cues code for different motion directions: do they give rise to discriminable patterns of neural signals, and is there evidence for their convergence onto a single ‘motion-in-depth’ pathway? To answer this, we use a decoding algorithm to test whether, and when, patterns of electroencephalogram (EEG) signals measured from across the full scalp, generated in response to CD- and IOVD-isolating stimuli moving towards or away in depth can be distinguished. We find that both MID cue type and 3D-motion direction can be decoded at different points in the EEG timecourse and that direction decoding cannot be accounted for by static disparity information. Remarkably, we find evidence for late processing convergence: IOVD motion direction can be decoded relatively late in the timecourse based on a decoder trained on CD stimuli, and vice versa. We conclude that early CD and IOVD direction decoding performance is dependent upon fundamentally different low-level stimulus features, but that later stages of decoding performance may be driven by a central, shared pathway that is agnostic to these features. Overall, these data are the first to show that neural responses to CD and IOVD cues that move towards and away in depth can be decoded from EEG signals, and that different aspects of MID-cues contribute to decoding performance at different points along the EEG timecourse.

---

To navigate the visual environment successfully, the primate visual system must interpret signals that indicate movement through three-dimensional (3D) space, or ‘motion-in-depth’ (MID). Stereoscopic vision offers two potential cues to MID (Cumming & Parker, 1994; Harris, Nefs, & Grafton, 2008; Rashbass & Westheimer, 1961; Regan, 1993). First, the change in disparity over time (CD) can be computed from the temporal derivative of interocular differences (disparities) in retinal position. In this case, position is computed first, and temporal differences are computed second: objects moving towards or away from the observer have opposite-signed CD signals. Alternatively, the interocular velocity difference (IOVD) can be derived by reversing the order of the computation above: retinal velocities are computed first and the interocular difference between these velocities also provides a signed indication of 3D object velocity. Although two separate retinal computations underlie the initial processing of CD and IOVD-cues, the extent to which these cues are processed via separate *cortical* pathways is still unknown.

At the retinal level, CD and IOVD depend on different aspects of low-level stimulus properties. However, at some point binocular integration is required to extract both CD (by computing the interocular disparity) and IOVD (by computing the interocular velocity) information. As there is very little binocular interaction in the LGN, this binocular stage is almost certainly located in visual cortex (Casagrande & Boyd, 1996). In the case of CD, the extraction of binocular disparity presumably occurs in primary visual cortex (V1), but the sites of binocular motion computations remain unclear. Recent functional magnetic resonance neuroimaging (fMRI) and psychophysical studies suggest that 3D-motion might be encoded downstream; areas in and around MT have been shown to respond to MID (Likova & Tyler, 2007; Rokers et al., 2009) and MT neurons are selective for 3D-motion directions (Czuba et al., 2011; Joo et al., 2016; Rokers et al., 2009, 2011). Likewise, fMRI studies have identified that these cues both activate motion selective area MT (or its human homologue) and nearby regions, while electrophysiological studies in macaque have shown that CD and IOVD stimuli both drive responses in macaque MT (Czuba et al., 2014; Héjja-Brichard et al., 2020; Likova & Tyler, 2007; Rokers et al., 2009; Sanada & DeAngelis, 2014). However, psychophysical measurements of CD and IOVD suggest that these mechanisms adapt independently (Joo et al., 2016; Sakano et al., 2012), have different speed sensitivities (Shioiri et al., 2008; Wardle & Alais, 2013), and engage systems with different spatial resolutions (Czuba et al., 2010). Thus, CD and IOVD appear to be processed via relatively independent pathways that provide input, and potentially converge, in MT (Joo et al., 2016). The extent to which both 3D-motion direction and MID pathways remain separate throughout cortical processing is not known. Further, it is unclear whether these pathways converge at later stages of processing.

These questions can be addressed using electroencephalography (EEG) and multivariate statistical techniques. To date, no study has used EEG to investigate MID processing, and further, there is a scarcity of research testing for the existence of 3D-motion direction sensitive cells. Unlike fMRI and behavioural methods, the high temporal resolution of EEG allows one to test for changes in the pattern of neural response *across the timecourse* of a relatively short analysis period, with the underlying assumption that later timepoints reflect later processing stages that occur higher up the visual processing hierarchy. This processing hierarchy is not necessarily mapped to the anatomical hierarchy; feedback to early visual areas may occur at relatively late stages of functional processing (Bullier, 2001; Lamme & Roelfsema, 2000) and regions outside of visual cortex. The high temporal resolution of EEG comes at the cost of poor spatial resolution; one limitation is that this method does not allow us to pinpoint the cortical regions that drive such neural responses. Moreover, novel machine learning decoding allows one to identify timepoints at which the pattern of electrical responses distributed across the full scalp differ between both MID cue type and 3D-motion direction. We employ this powerful technique to investigate the independence of MID pathways in the cortex. We asked a series of questions about the *overall pattern* of neural signals generated by CD and IOVD stimuli moving in different 3D-motion directions and answered them using both within-cue and cross-trained decoding.

Here, we used EEG to measure signals from 64 channels across the scalp, generated in response to well-isolated CD and IOVD cues that either moved towards or away in depth. First, we asked if different 3D-motion directions give rise to different EEG responses by testing whether a multivariate pattern classifier (i.e. a decoder) could use the pattern of EEG signals to accurately discriminate motion direction (towards vs away) for both CD and IOVD cues. The ability to do this would suggest that different motion directions drive different neural populations, or alternatively, drive similar populations with differences in timing, synchrony, and coherence. One confound here is that CD stimuli necessarily have a start and end position in 3D space. To control for this, our CD were designed to maintain the same *average* depth across a stimulus cycle irrespective of 3D-motion direction, but this meant that they started from different sides of the horopter. In theory therefore, our CD motion direction decoding could be based on the time-averaged disparity around the stimulus onset or offset. To examine this, we ran a second experiment that used dynamically updating static CD stimuli (‘static disparity stimuli’) that were located either near or far in depth, with no stereomotion information. Thus, we also asked whether we could decode relative stimulus position of these static disparity stimuli (near vs far).

In addition, we asked if CD and IOVD cues generate different response patterns. Although our stimuli were matched as far as possible in their low-level properties (dot density, size, and contrast), they necessarily differed in some important ways (e.g. dot duration and velocity). We hypothesised that CD and IOVD stimuli would drive different neural populations because of these differences and, potentially, because they may also be processed along anatomically independent pathways. To address this, we tested whether, and when, the decoder could use EEG signals to accurately discriminate between MID cue types (CD vs IOVD).

CD and IOVD stimuli rely on fundamentally different low-level features, thus, if cross-trained decoding is possible at some point in the EEG timecourse, it would indicate some contribution of shared MID information, perhaps due to a mechanism that is agnostic to such features. Therefore, our final question asked whether MID cue decoding performance relied on unique CD and IOVD signals, or if there was shared information between these two signals that would, for example, allow us to decode CD motion direction based on IOVD motion direction data. To test this, we cross-trained the decoder and examined whether it could accurately discriminate CD motion direction cues after being trained using only IOVD motion direction cues, and vice versa.

## Methods

### Participants

In the first experiment, twelve healthy participants (mean age = 24 years, eight males) were recruited from the University of York. All participants completed a pre-screening to ensure they had functional stereoscopic vision. Two participants did not pass the prescreening and did not participate in the experiment. Thus, ten participants (eight males) completed a one-hour EEG session. In a second ‘static disparity’ experiment, nine participants (mean age = 27 years, four males, including three participants from the previous experiment) completed a one-hour EEG session. All participants had normal or corrected-to-normal vision and provided written consent before participating in the study. Experiments were conducted in accordance with the Declaration of Helsinki and the experimental protocol was approved by the ethics committee at the Department of Psychology, University of York.

### Stimulus display

Stimuli were presented on a VIEWpixx/3D (VPixx Technologies, Saint-Bruno, Quebec, Canada) display (1920 x 1200 pixels,120 Hz refresh rate). The display had a mean luminance of 100 candela/m^2^ and was gamma corrected using a photospectrometer (Jaz; Ocean Optics, Largo, FL, USA). Participants viewed the display from 57 cm and were required to wear wireless liquid crystal display (LCD) shutter googles that were controlled by an infrared emitter (NVIDIA GeForce 3D; NVIDIA, Santa Clara, CA, USA). Here, binocular separation of the stimuli, with minimal crosstalk, was achieved by synchronising the VIEWpixx refresh rate (120 Hz, thus 60 Hz per eye) with the toggling of the LCD shutter goggles (Baker et al., 2016).

### Stimuli

#### CD and IOVD Stimuli

The stimuli were similar to those used in previous papers (Kaestner et al., 2019; Maloney et al., 2018). Briefly, MID stimuli consisted of random dot patterns that were designed to experimentally dissociate CD- or IOVD-based mechanisms. The low-level properties of the dots were matched between cue types as far as possible. In all conditions, the dots were anti-aliased and had a Gaussian profile (σ= 0.05°). The black and white dots were positioned at a density of one dot per 1° of visual angle^2^ on a mean luminance grey background, with a Michelson contrast of 100% (50:50 black:white). Stimuli were presented using MATLAB 2014a (The Mathworks Inc., Massachusetts, USA) and the Psychophysics Toolbox Version 3 (Brainard, 1997).

For CD stimuli, MID information is carried by changes in the disparities of pairs of dots presented to the left and right eye. To remove any velocity cues, dots can be replaced at new random positions on each video refresh. Thus, CD stimuli consisted of temporally uncorrelated dynamic random dot patterns, where each frame consisted of a set of binocularly correlated, randomly positioned dot patterns, only differing in a lateral shift in retinal disparity between the left and right eye (see **Figure 1A**). CD stimuli changed unidirectionally in disparity (i.e. the stimulus moved either towards or away in depth). This shift in disparity followed a single linear ramp that began at the far point and finished at the near point for towards motion, and the opposite points for away motion. The mean stimulus-averaged depth relative to fixation for both stimuli was therefore zero. The near and far points were identical for each with a peak binocular disparity of ±32 arcmin which is below the fusion limit for typical human observers (Howard & Rogers, 2002; Norcia & Tyler, 1984).

**Figure 1.**
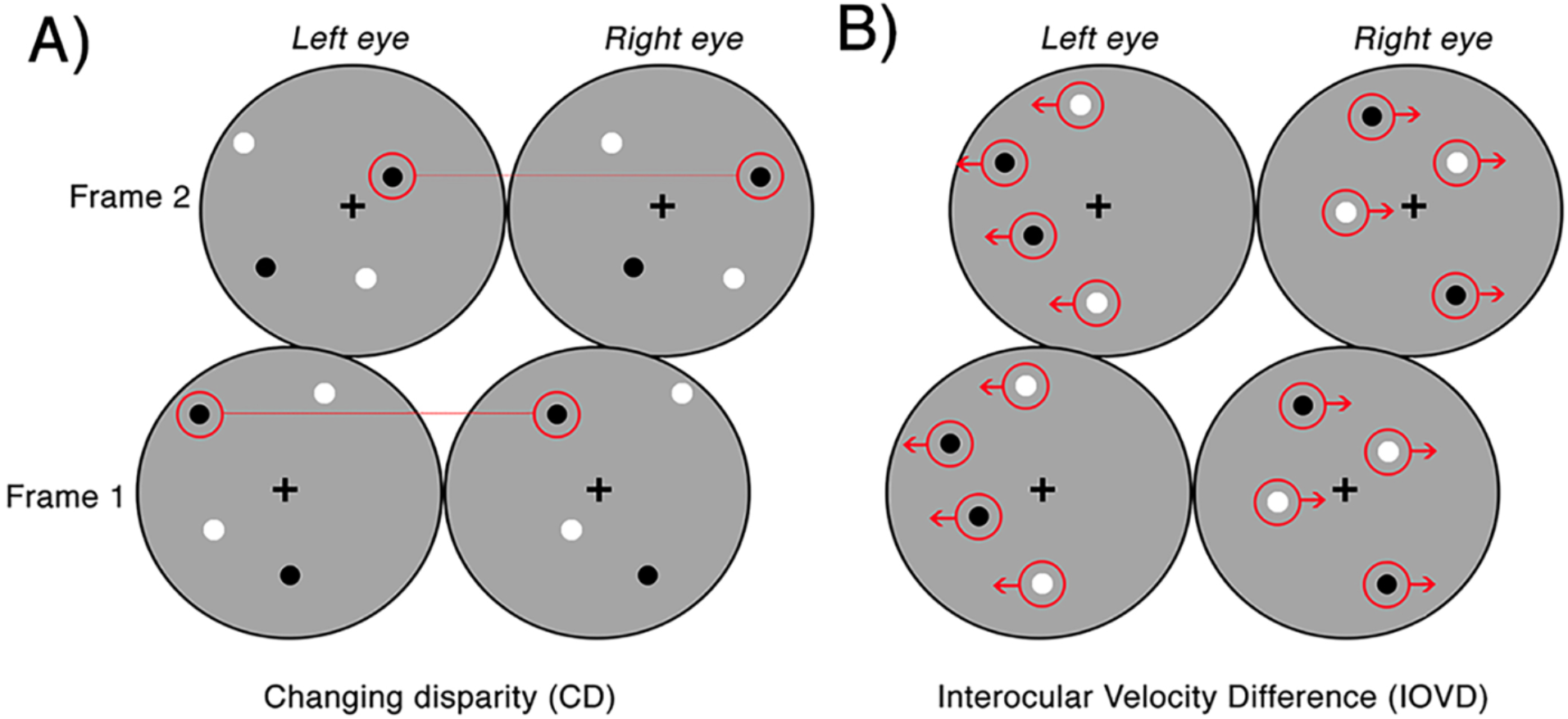
Examples of the CD and IOVD stimuli presented across two consecutive frames. Panel A) illustrates CD, where the dots are positioned randomly and paired between the left and right eyes and only differ in their lateral shift in disparity across each video frame. Panel B) illustrates IOVD, where the dots laterally shift through time and there is no pairing of dots between the left and right eyes.

For IOVD stimuli, MID information is carried by the relative interocular velocities of retinal features across time. No fine-grained interocular matches are required (or indeed possible). Our IOVD stimuli consisted of fields of moving dots with the left and right eyes seeing uncorrelated dot patterns. The patterns moved coherently across time and the velocity of this motion was equal and opposite in the two eyes (see **Figure 1B**). The lateral shift of the dots that defined the unidirectional MID was, again, in the form of a linear ramp. To maintain uniform dot density and equal visual transients across the stimulus, equal numbers of dots reached the end of their lifetime (50ms) and were ‘reborn’ in a new position on the stimulus for each frame of the stimulus. For IOVD stimuli, the maximum monocular lateral displacement was ±128 arcmin.

Thus, to ensure that no disparity information leaked into IOVD stimuli, the spatial positioning of the IOVD dot patterns was organised so that they fell into two horizontal strips that were two dot widths wide (~0.5°). The strips alternated across the eyes to ensure that they did not coincide on the two retinae (Shioiri et al., 2000, 2008). On the rare occasion when a dot fell near the border of a horizontal strip in the left eye that was close to the dot near the border of the horizontal strip in the right eye, these dots were assigned opposite contrast polarity to disrupt any potential disparity signals (Maloney et al., 2018).

All stimuli were presented within an annulus that had an inner radius of 1° and an outer radius of 6° (see **Figure 2**). The contrast of the dots appearing at the edge of the annulus was smoothed with a cosine ramp that was 0.5° wide. To provide relative depth information, a circular central fixation lock (0.4° radius) surrounded the fixation cross, while a circular outer fixation lock (11.75° radius) surrounded the edge of the display. These locks were split into black and white quadrants that served as a set of nonius lines to assist in gaze stabilisation and the fusion of the images presented to the two retinae. An example of a single frame from the stimulus is presented in **Figure 2**. Animated examples of CD and IOVD stimuli are available in Supplementary Materials of previous work (Kaestner et al., 2019).

**Figure 2.**
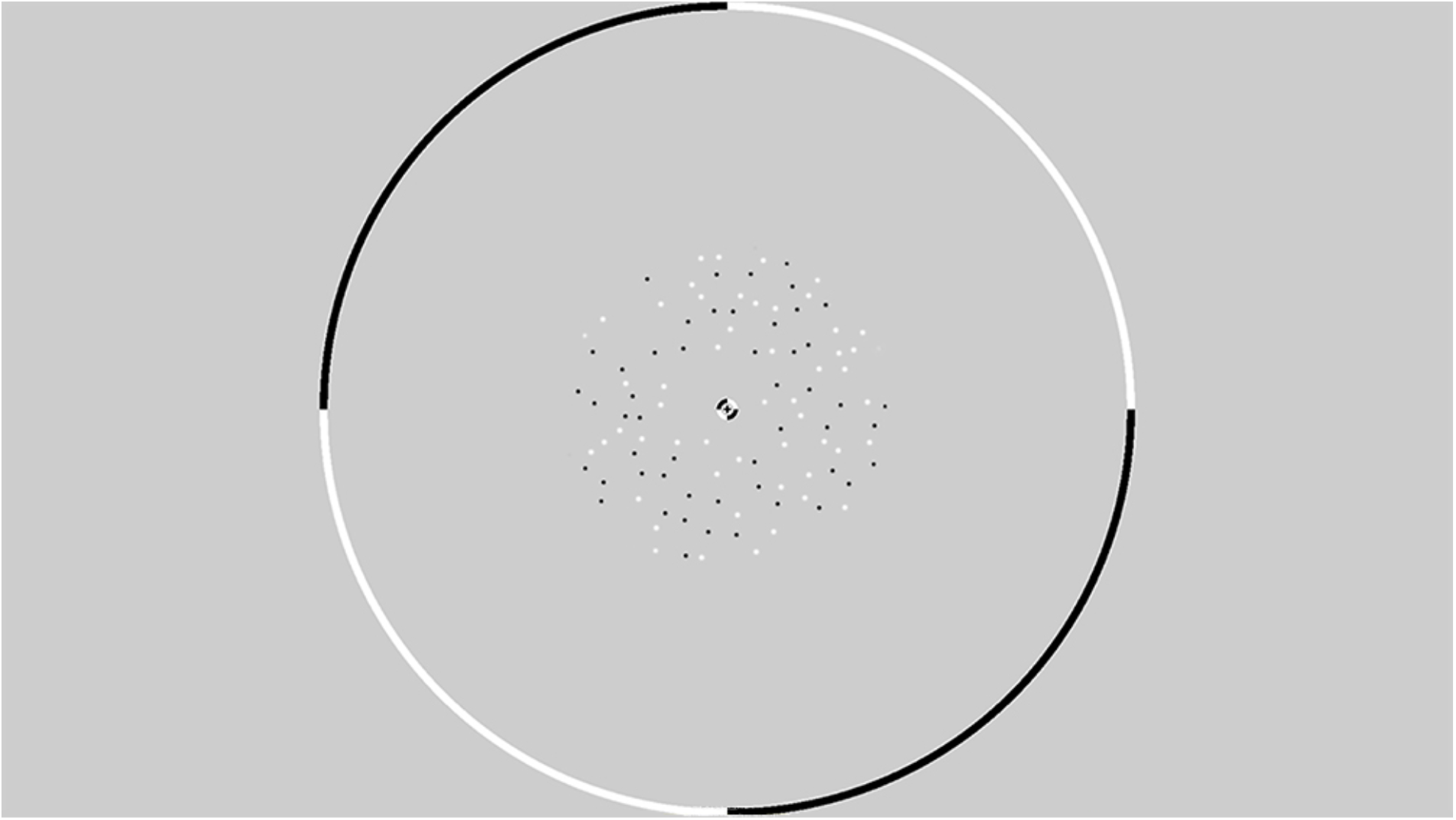
An example from a single frame of the random dot stereogram pattern that is presented within a circular fixation lock. The dots are presented in an annulus with an inner radius of 1° and an outer radius of 6°.

#### Static disparity stimuli

Using fixed far and near endpoints for CD stimuli means that a decoder discriminating between CD stimuli moving towards and away in depth could potentially rely on static disparity information at either the beginning or the end of the stimulus presentation, rather than the time-averaged MID information. We ran a control experiment that examined whether the decoder could distinguish between dynamically updating but depth-fixed CD stimuli located at either the near or far positions in depth. The near and far static positions in depth were identical to the extremes of depth at the start and end points of the MID CD stimuli. These extreme positions were chosen as they would deliver the largest static disparities available to us, thus producing the largest signal. These disparity stimuli had identical durations (250 ms) and dot update rates to the CD stimuli, however the stereomotion velocity was set to zero.

### Experimental design

Participants’ 3D vision was tested using the TNO Stereo Test, 19^th^ edition (Lameris Ootech, NL) to ensure that they had functional stereo-acuity (with thresholds at or below 120 arcsec). This test screens for static depth perception (rather than MID perception) as traditionally defined in clinical settings (see Maloney et al., 2018). During the EEG cap set-up, participants were presented with a demonstration of each of the four stimulus conditions on the VIEWpixx display, whereby the stimuli oscillated towards and away continuously according to the linear ramp. This allowed participants to practice identifying towards and away motion directions and CD and IOVD cues. These demonstrations did not require a keyboard response and were presented until the participant felt that they were confident in identifying all four stimulus conditions and ensuring that they could perceive stereomotion. Following this, participants completed two practice runs of the experiment itself. During testing, participants could return to practice runs if necessary.

Participants completed six blocks of 220 stimulus trials. Within each block the four stimulus conditions (CD towards, CD away, IOVD towards, and IOVD away) were randomised and occurred with equal frequency. For each participant, we retained 210 of these trials to account for any potentially dropped triggers during testing. In the static disparity control experiment, participants completed three blocks of 220 stimulus trials, with 210 retained, and each block contained two stimulus conditions, ‘CD near’ or ‘CD far’, that were also randomised and occurred with equal frequency. Participants were given a one-minute break between each block. For each trial, the stimulus probe had a duration of 250 ms, which corresponded to one full linear ramp (i.e. the stimulus followed a unidirectional trajectory over the 250 ms duration). The stimulus probe was followed by an inter-trial interval (ITI). The length of the ITI was selected randomly from a uniform distribution (lower bound = 1 s, upper bound = 3 s). Participants were instructed to fixate on the central cross throughout the entire experiment. While fixating, participants completed a task in which they were required to respond, using their right hand, to indicate whether the stimulus was moving towards (2 on the number pad) or away (8 on the number pad) from them. The keyboard response was recorded, and participant accuracy is available in Supplementary Materials (**Figure S1**). For the static disparity experiment, participants were instructed to indicate whether they perceived the stimulus as being near (‘2’) or far (‘8’) in depth. Participants received audio feedback in the form of a tone to indicate incorrect responses only and were not shown their overall response accuracy.

### EEG recording and collection

EEG data were collected at 1 KHz from 64 electrodes that were distributed across the scalp according to the 10/20 system in an ANT WaveGuard EEG cap and digitized using the ASA Software (ANT Neuro, Hengelo, NL). The ground electrode was placed posterior to electrode *FPz* and the signal from each channel was referenced to a whole-head average. Eye-blinks were recorded using two vertical electrooculogram electrodes. Individual records were inspected for excessive blinking and we found that this was not an issue with our data. These signals were amplified, and the onset of each stimulus trial was recorded in the EEG trace using a low-latency digital trigger. Data were exported to MATLAB 2018a for offline analysis using customised scripts.

### EEG pre-processing and bootstrapping

For each participant, EEG data were epoched into 1000 ms epochs at −200 ms to 800 ms, relative to stimulus onset at 0 ms (see **Figure 3**). A low-pass filter of 30Hz was applied to remove high frequency noise, including line noise. Each epoch was then down sampled from 1000 Hz to 125 Hz via linear interpolation. Thus, we retained a sampling pool of 1260 EEG epochs (210 trials x 6 blocks) that included each of the four stimulus conditions. To gain an estimate of the variance of our classification results we used permutation testing. For each stimulus condition, we bootstrapped the data of 10 epochs from the sampling pool. These 10 epochs were averaged together to create a mean bootstrapped epoch. This was repeated for each stimulus condition. For each participant, this process was repeated 21 times. Thus, for each participant, and each stimulus condition, there were 21 mean bootstrapped EEG epochs, for each of the 64 electrodes. Bootstrap resampling and averaging were done in parallel across all EEG channels. Data were then z-scored to standardize across electrodes. This process was repeated for 1000 iterations of a bootstrapped support vector machine (SVM) classifier, with each iteration generating a unique set of 21 x 21 mean bootstrapped EEG epochs for each participant, by averaging together new sets of sample epochs from the sampling pool. This generated unique input data for each iteration of the SVM.

**Figure 3.**
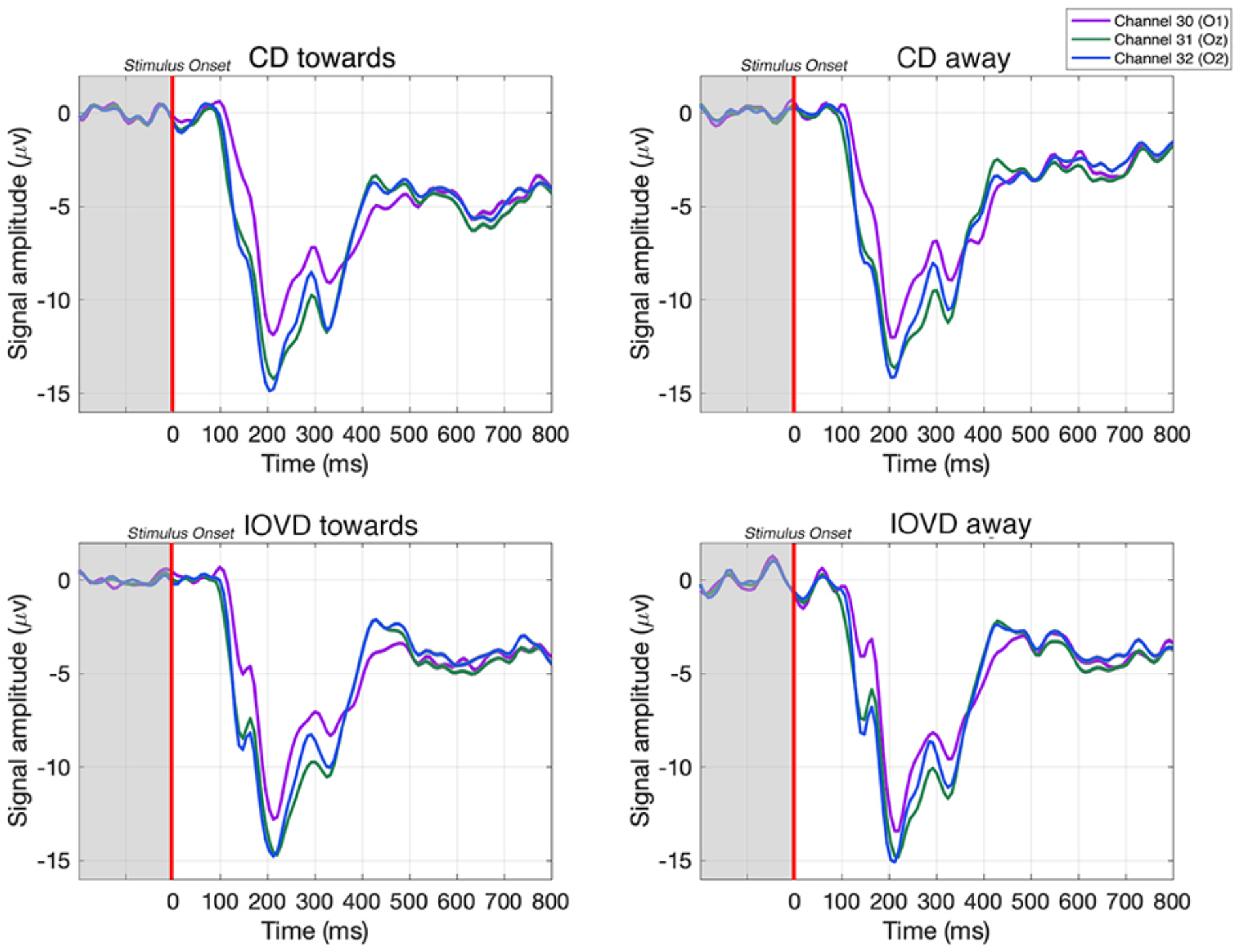
Examples of averaged EEG waveforms from a single participant, taken from 3 occipital electrodes in response to each stimulus type. Waveforms are averaged over 210 trials. Each epoch is −200 to 800 ms and the stimulus is presented between 0 – 250 ms. A relative change in EEG signal begins around 100 ms after stimulus onset.

### Bootstrapping keyboard response and percept control analysis

In a supplementary control analysis, we asked whether our results could be driven by motor signals (due to the button press – ‘2’ or ‘8’ on the keypad) rather than visual inputs. To check this, we asked whether we could decode EEG data that had been pooled into keyboard response conditions (with balanced stimulus types), rather than stimulus conditions. As we decoded using all the electrodes across the scalp, our goal was to test whether motor responses (i.e. responses that occur due to using the middle or index finger on the same hand to indicate if the stimulus was moving towards or away) had any influence on decoding performance (Cottereau et al., 2011, 2013, 2014). Thus, if our SVM is inaccurate at decoding keyboard-response, it would indicate that the EEG signals associated with the keyboard response have little influence on decoding accuracy, confirming that our overall decoding results are driven predominately by sensory inputs. The methods and results for this control analysis can be found in *Supplementary Materials: Decoding keyboard responses: motor responses do not contribute to decoding performance* (**Figure S2**). Further, we used these keyboard responses to run an additional supplementary analysis that tested whether the *percept* of MID direction could be decoded from the EEG signals. Here, we used each participants’ keyboard responses to create mean bootstrapped EEG epochs based on whether a participant *perceived* the stimulus moving towards or away in depth, irrespective of the actual direction, or cue, of the stimulus. A description of the analysis and the results are presented in detail in *Supplementary Materials: Decoding the directional perception of MID based on behavioural responses* (**Figure S3**).

### Statistical analysis: Decoding EEG signals using a support vector machine (SVM)

To decode MID information from our EEG data, we bootstrapped a pairwise SVM classifier with a linear kernel in MATLAB 2018a using the LIBSVM toolbox, v. 3.23 (Chang & Lin, 2011). This allowed us to run pairwise decoding at each of the 125 time-points, thus obtaining decoding accuracies across the EEG time-course. In the case of our EEG data, decoding can only occur at above-chance levels if different stimuli produce patterns of electrical activity at the scalp that differ in some consistent manner on an individual basis. For each participant, decoding was performed at each of the 125 time points (from −200 ms to 800 ms), using data from all 64 electrodes. The advantage of this decoding technique is that it does not assume consistency in the pattern of electrical activity across participants, but rather, the decoding is run on individuals. Examples of the pattern of EEG response across the scalp at early-, mid-, and late-stages of the EEG time course are available in *Supplementary Materials: Scalp distributions* **Figure S4** and **Figure S5**. Each electrode had equal weighting in its contribution to decoding accuracy. EEG data were bootstrapped repeatedly (as described above) and run through 1000 iterations of the SVM to derive a mean decoding accuracy at each time point, for each participant. Significant decoding accuracy was assessed across time. Here, decoding accuracy was deemed significantly different from chance (50%) using a non-parametric cluster corrected *t*-test (as described below).

Decoding accuracy was estimated using a leave-one-out classification procedure. On each run, the SVM is trained on all but one of the EEG epochs. This left-out EEG epoch serves as the test data that will be classified by the SVM. We trained the SVM on 40 EEG epochs while 1 EEG epoch was left-out as the test data (Boser et al., 1992; Cortes & Vapnik, 1995; Wang, 2018). Thus, the SVM was trained to decode between two stimuli by learning to discriminate between the pattern of EEG responses arising from a pair of stimulus conditions at a single time-point. The SVM then predicted the class of the left-out test EEG epoch (Grootswagers et al., 2017). If the SVM can accurately classify the test EEG epoch into its correct class, decoding accuracy will be high. If the SVM cannot classify the test EEG epoch into its correct class, decoding accuracy will fall around chance; this would suggest that there are no meaningful differences in the EEG response elicited by the two stimulus conditions in that comparison. A decoding accuracy is obtained for each of the 125 time-points across the EEG time-course. Chance baselines were verified by shuffling class labels. This confirmed that SVM performance was not driven by artefacts in the signal processing pipeline (see Supplementary Materials **Figure S6** and **S7** for results of decoding accuracies after shuffling labels).

Additionally, we ‘cross-trained’ the SVM classifier to test if shared information between CD and IOVD-cues could be used to decode 3D-motion direction. This analysis differed from that above. Here, the SVM is trained to decode 3D-motion direction for a single MID cue-type (i.e. CD towards vs CD away) and *outputs a classification model.* The decoder then uses this model to decode new, unseen data that comes from a *different pair of conditions* (i.e. IOVD towards vs IOVD away). Thus, in the cross-trained analysis, we trained the SVM to decode CD motion direction and used the output model to decode IOVD motion direction, and vice versa.

### Non-parametric Cluster Correction

To test for significance, decoding accuracy was compared to a chance baseline (50% decoding accuracy) using a non-parametric cluster comparison. By definition, the time points at which the decoder performs at above-chance (as assessed from the distribution of bootstrapped SVM classification accuracies) must have an overall pattern of EEG signals that differs significantly between stimulus conditions. However, the large number of time points (125) can inflate the false discovery rate due to multiple comparisons.

To solve this, we used a non-parametric cluster correction test. This method is commonly used in EEG and takes advantage of temporally clustered significant time-points to compute the *t*-statistic (Maris & Oostenveld, 2007). The approach involves the following: first, for each time-point, decoding accuracy is compared to 50% chance baseline by means of a one-sample *t*-test. To reduce the number of multiple comparisons, these significant time-points are then clustered based upon their temporal adjacency and commonality. Next, cluster level *t*-values are calculated by summing the *t*-values within each cluster. A null distribution is computed via a permutation test, and we ask whether the cluster-level *t*-value is greater than one that would occur by chance (using this null distribution). Any clusters with a significant pooled *t*-value, when compared to the null distribution, are identified. If the *p*-value for a cluster is smaller than the cluster threshold (*p* <.05), it is deemed to be a significant cluster.

## Results

### EEG Results

All our multivariate decoding was performed using a series of binary comparisons. First, we tested if we could discriminate between towards and away 3D-motion direction responses when pooled across MID cue type. Next, we tested if we could discriminate between CD and IOVD cue type responses when pooled across 3D-motion directions. Following this, we asked if we could decode stimulus 3D-motion direction responses *within* MID cue type, and between MID cue type responses *within* stimulus 3D-motion direction; therefore, comparing between CD towards against CD away cues, IOVD towards against IOVD away cues, CD towards against IOVD towards cues, and CD away against IOVD away cues. Next, we examined the reliance of CD decoding performance on static disparity information by testing whether we could discriminate responses from static disparity stimuli at either near or far positions in depth. Finally, we cross-trained the decoder and tested if it could distinguish between CD towards against CD away cues using a decoding model that was trained using IOVD motion direction data, and vice versa.

Here, we present decoding accuracy for these binary comparisons. We report decoding accuracy across time (from stimulus onset at −200ms through to 800ms). Likewise, we report whether this decoding accuracy was significantly above a chance baseline of 50% decoding accuracy (*p* < .05), across time.

### Decoding between 3D-motion direction and cue type: MID cues moving towards and away in depth give rise to independent neural responses

First, we tested whether the decoder could discriminate between EEG signals generated in response to towards and away 3D-motion directions when the data was pooled across MID cue type. As illustrated in **Figure 4** (red line), after collapsing over CD and IOVD cue types, we were able to decode towards vs away 3D-motion direction at above-chance levels from 320 ms until 632 ms after stimulus onset, with accuracy peaking at 65% at 536 ms (*p* < .05). Thus, our results show that we could decode the direction of motion through depth (i.e. either moving towards or away), independent of cue type.

**Figure 4.**
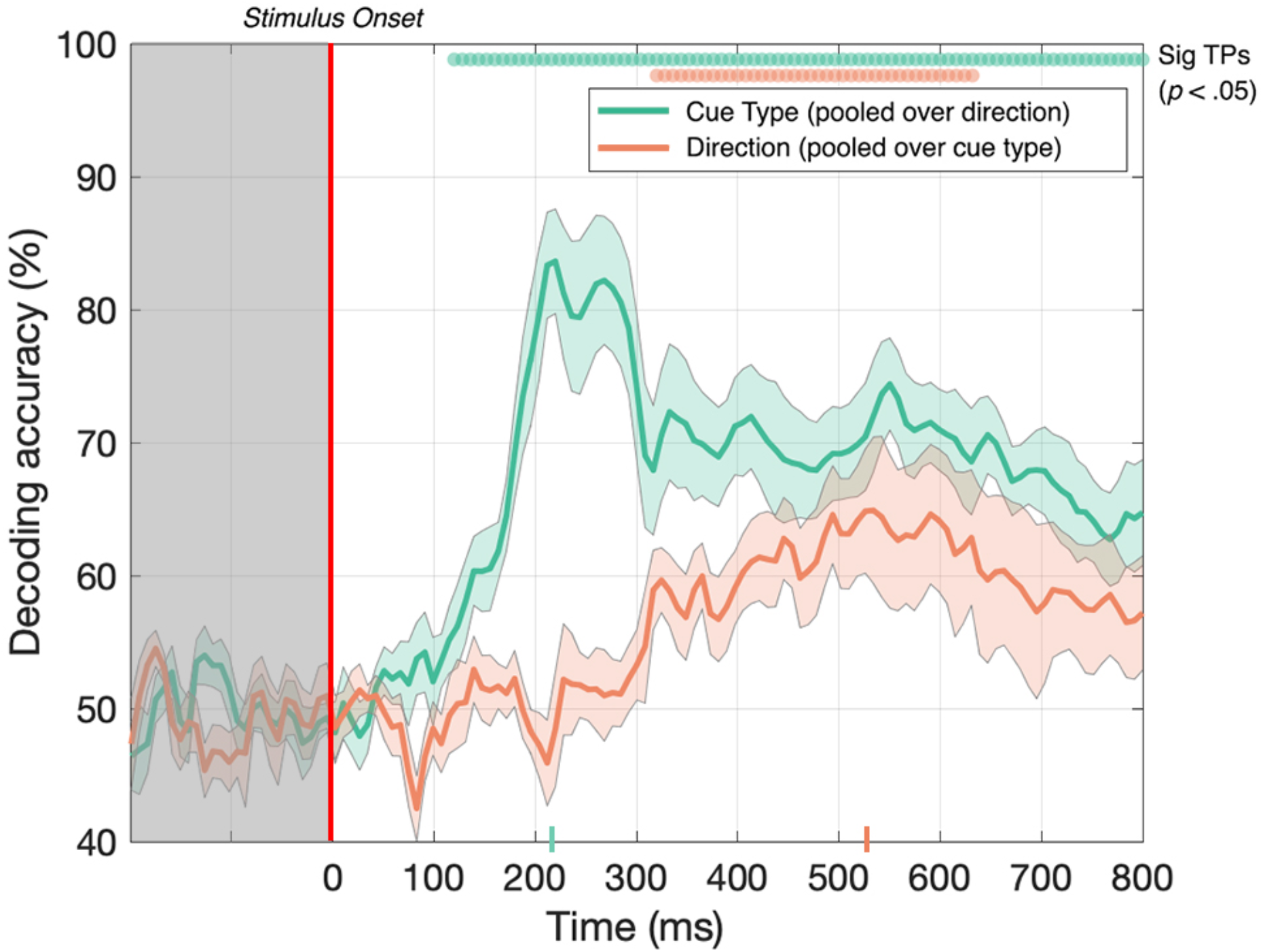
Decoding accuracy across 125 time points for cue type (pooled over motion direction) in green and 3D-motion direction (pooled over cue type) in red. The coloured ticks on the x-axis represent the time point of peak accuracy for each condition. Green and red circles indicate the points of time when the Bonferroni-corrected t-tests were significant (p < .05). Shaded error bars represent ±1SE of the bootstrapped mean (1000 iterations).

Next, we tested if the decoder could discriminate between CD and IOVD cue responses when the data was pooled across 3D-motion direction. As illustrated in **Figure 4** (green line), EEG responses to CD and IOVD cues (irrespective of 3D-motion direction) could be decoded relatively early in the EEG timecourse. Decoding occurred above chance at 120 ms after stimulus onset and continued at above chance levels across the remaining timecourse (*p* < .05). Here decoding performance peaked at 224 ms and reached 84% accuracy. The decoding performance for IOVD vs CD at 120 ms is the earliest difference that we see in this study. Notably, the peak of decoding accuracy occurred much earlier than the peak for decoding motion direction (towards vs away).

We hypothesise that the ability to decode between CD and IOVD cues is due to the low-level properties of these cues themselves. Although we matched the low-level cue properties of the CD and IOVD stimuli as far as possible, they inevitably differed in some respects. CD stimuli (deliberately) contained no coherent monocular motion energy while the dots in the IOVD stimuli travelled short distances on the retina before being refreshed. These differences are intrinsic to the cues themselves – a MID system that isolates CD will necessarily rely on different low-level aspects of the stimulus than an IOVD pathway, and these differences in stimulus properties cannot be avoided, although that is not to say that these cues do not drive the unique responses seen here. Additionally, in principle, differences in the cortical signals generated by different eye-movements patterns could also contribute to CD vs IOVD decoding performance. Moreover, the later peak for 3D-motion direction decoding may be due to increased time for the integration of MID information and, potentially, the role of feedback from higher level brain mechanisms when decoding between 3D-depth directions.

### Decoding within motion direction and cue type

Decoding performance *pooled* across stimulus 3D-motion direction and cue type tells us little about the computations underlying CD and IOVD cues. For example, IOVD and CD may recruit entirely different neural populations, or alternatively, these cues may activate similar neural populations but generate unique responses that differ in their timing, coherence, or synchrony. Because of the way that we pooled the data (i.e. pooled over MID cue type, and pooled over 3D-motion direction), it is possible that decoding performance is driven by only *one* of the two pooled elements at any particular time point. For example, CD 3D-motion direction decoding performance may peak at early stages of the timecourse, and IOVD 3D-motion direction may peak at later stages of the timecourse. To gain a more complete understanding of the neural responses driven by each cue type and 3D-motion direction, we split our data into individual conditions and ran a series of further comparisons.

Here, we asked whether we could decode 3D-motion direction *within* individual cue type and whether we could decode cue type *within* individual 3D-motion directions. Our data (**Figure 5**) show that we could do so at above-chance levels for all comparisons. The decoder obtained above-chance performance in decoding the 3D-motion direction of both CD and IOVD stimuli (*p* < .05). CD towards vs CD away stimuli could be decoded from 320 ms - 672 ms, then again from 688 ms – 800 ms, with a relatively late peak in decoding accuracy of 68% at 592 ms (**Figure 5**, red line). IOVD towards against IOVD away stimuli could be intermittently decoded from 296 ms, with a decoding peak of 58% at 536 ms (**Figure 5**, light blue line). The decoder could first distinguish between IOVD direction 24 ms before CD direction, and IOVD direction decoding accuracy (536 ms) peaked 56 ms before CD direction decoding accuracy (592 ms).

**Figure 5.**
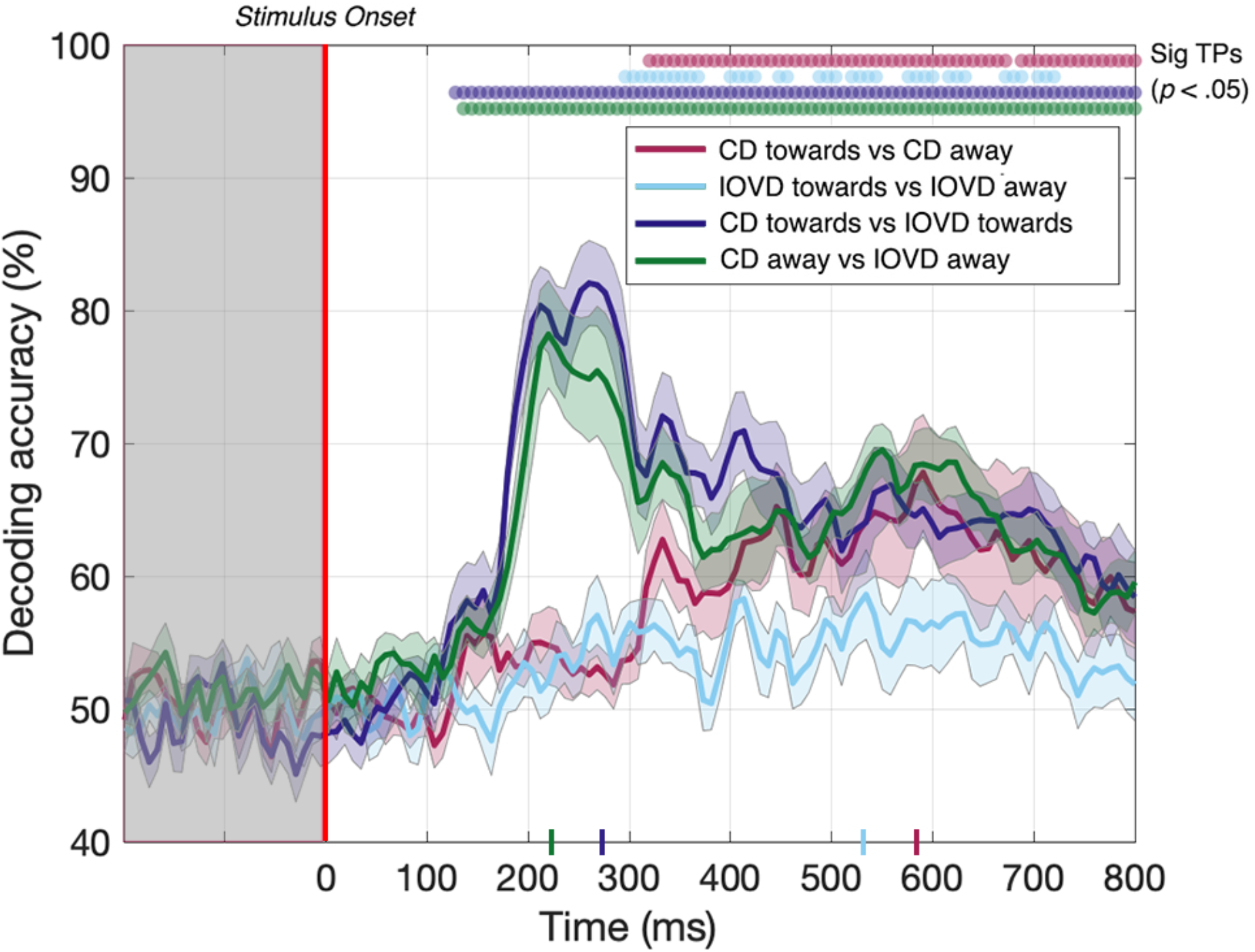
Pairwise decoding accuracy across 125 time points for four stimulus conditions. Red, light blue, dark blue, and green dots indicate time points when the Bonferroni-corrected t-tests were significant for each condition (p < .05) and the coloured ticks on the x-axis represent the time point of peak decoding performance for each stimulus condition. Shaded error bars represent ±1SE of the bootstrapped mean (1000 iterations).

The ability to resolve EEG responses to 3D-motion direction is intriguing. The ability of the decoder to discriminate between towards and away 3D-motion direction shows that differences in response can be resolved using EEG, and that these responses are relatively independent from each other in the cortex. One might hypothesize that this independence is due to an intermingled network of neurons that are selective for different 3D-motion directions.

For completeness, we also tested if the decoder could discriminate between CD and IOVD cues *within* each 3D-motion direction. Decoding between CD towards and IOVD towards stimuli (**Figure 5**, dark blue line) was significant from 128 ms and peaked at 82% accuracy at 264 ms (*p* < .05). Next, decoding between CD away and IOVD away stimuli (**Figure 5**, green line) was significant from 136 ms and accuracy peaked at 78% at 224 ms (*p* < .05). Thus, significant accuracy in decoding between cue-type within individual motion directions begins soon after stimulus onset. Decoding performance accuracy remains high along the EEG timecourse, regardless of motion direction; an effect that appears to be driven by unique neural responses to the fundamentally different low-level properties of CD and IOVD cues. Extended results of decoding between CD towards against IOVD away, and IOVD towards against CD away, are available in Supplementary Materials **Figure S8**.

### Decoding static disparity stimuli: Static depth information contributes to early stages of CD direction decoding, but not late stages

To ensure the same average stimulus disparity over the course of each trial, our CD stimuli began at one extreme in depth (i.e. either near or far) before traversing towards or away from the participant. However, the instantaneous disparities at the start and end of each presentation are therefore correlated with the motion direction, and at least part of the decoding performance for the CD towards vs CD away condition may potentially be driven by this static depth cue, rather than MID information. This confound is not present for the IOVD condition because these stimuli do not have a well-defined 3D location. To measure the extent to which our CD motion direction decoding was indeed due to MID information, rather than static depth information, we ran a second experiment to test if the decoder could discriminate between dynamically-updating static disparity stimuli that were held at one of two fixed depth locations – near or far. The static disparity stimuli used in this control were more informative about a single 3D depth than those used in the MID condition (because they signalled the same depth for a longer period). Thus, we hypothesize that the increase in decoding performance in the MID condition is due to 3Dmotion rather than the static depth confound. Conversely, static disparity information may have some contribution to MID CD decoding performance at time points where near and far static disparity stimuli can be decoded.

In the static disparity control condition, we identified a short set of time points where the decoder could distinguish between near and far depths. Our data showed that decoding was possible between 288 – 312 ms (**Figure 6**, green line). Thus, the decoder *could* distinguish between static disparity stimuli located near or far in depth for a short ‘early’ period around 300 ms. However, this differs from the MID CD condition (**Figure 6**, red line - identical timecourse from **Figure 5**), where CD towards vs CD away decoding was similarly possible from around 300 ms but *continued* to be significant across the rest of the EEG timecourse. Thus, transient static depth information may contribute to the earliest stages of MID depth decoding, however, the subsequent stages of decoding must be attributed to the smooth motion information that was not present in these stimuli.

**Figure 6.**
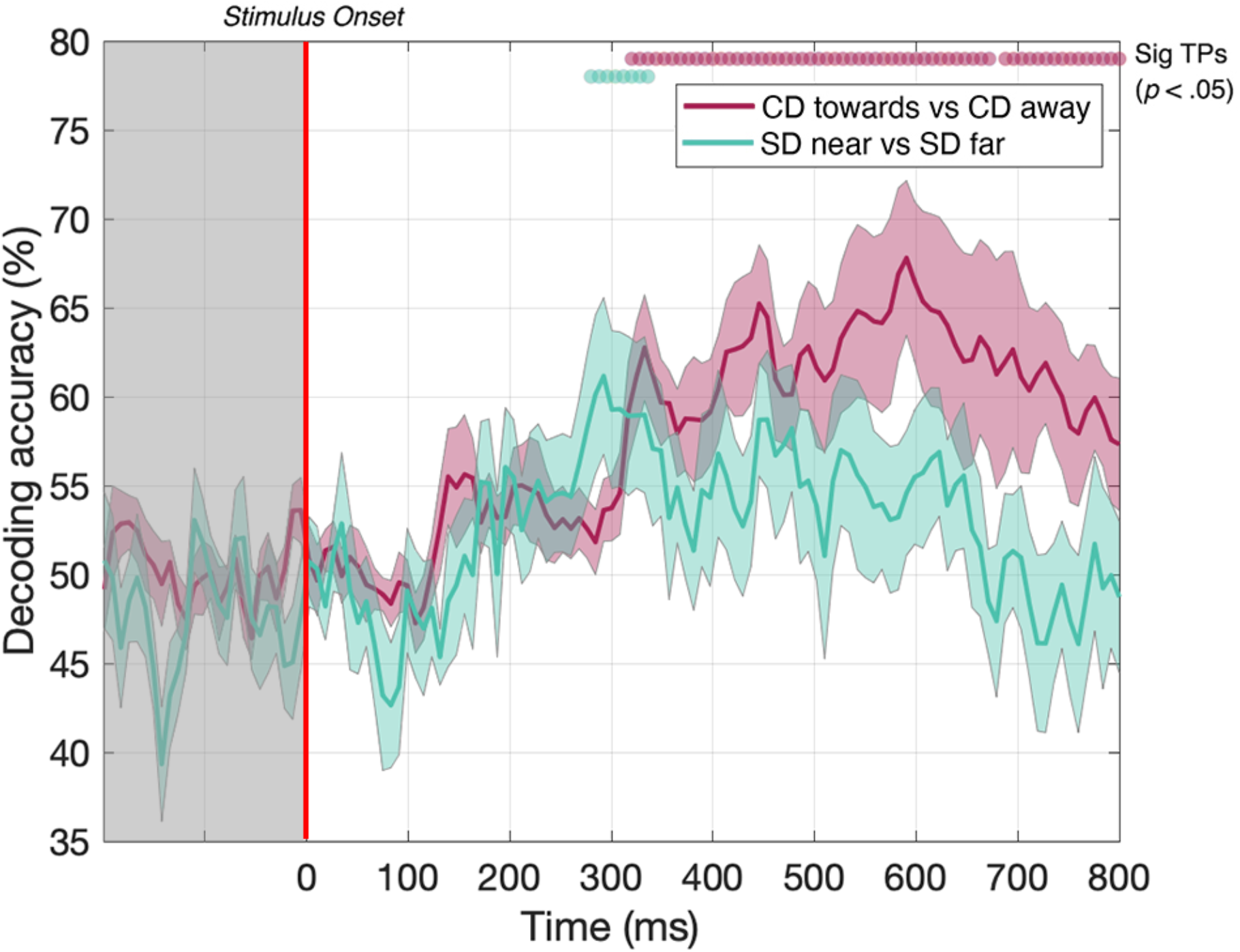
Pairwise decoding accuracy across 125 time points for static disparity (SD) stimuli that are presented either near or far in depth, and for CD stimuli moving either towards or away in depth. Red and green dots indicate time points when the Bonferroni-corrected t-tests were significant (p < .05). Shaded error bars represent ±1SE of the bootstrapped mean (1000 iterations).

### Decoding from a cross-trained decoder: early performance is associated with cue differences while later performance might extend from a shared MID pathway

Next, we examined whether CD and IOVD signals are entirely independent, or if neural information about stimulus motion direction is shared between these mechanisms at some point. To examine this, we ‘cross-trained’ the decoder. Here, we trained the decoder using towards vs away CD responses and then asked whether this CD-trained decoder could distinguish responses to different directions of IOVD motion, and vice versa. If CD and IOVD pathways are independent from each other, one could expect decoding accuracy to be close to 50% (chance), as a decoder trained on signals from one cue (i.e. CD) should not be helpful in decoding signals from the other cue (i.e. IOVD). However, if cross-trained decoding is possible along the EEG timecourse, it would suggest some cross over between CD and IOVD pathways, with the contribution of a mechanism that is agnostic to whether CD or IOVD cues deliver the MID. Further, because IOVD stimuli have no unique depth, this cross-validation information must be independent of confounds from pure disparity cues.

Remarkably, we could decode 3D-motion direction from the cross-trained classifier for significant periods at a latency of approximately 500 ms. Specifically, IOVD motion direction could first be decoded from a CD-trained classifier at 512 ms after stimulus onset with 56% accuracy (*p* < .05), with intermittent significance through to 800 ms (see **Figure 7**, purple line). A similar result was found for decoding CD motion direction from an IOVD-trained classifier: significant decoding first occurred at 488 ms after stimulus onset and again at 56% accuracy (*p* < .05), through to 552 ms, and was then intermittently significant through to 648 ms (see **Figure 7**, green line). Importantly, decoding was not possible during early stages of the stimulus presentation period (i.e. before 250 ms) or soon thereafter. The ability of the cross-trained decoder to perform 3D-motion direction decoding from a classifier trained on the opposing cue type indicates similarities in the pattern of response at later stages of MID processing. One possibility is that the earlier classification performance we see in our original CD vs IOVD decoding (**Figures 4 and 5**) is due to low-level stimulus differences in CD and IOVD cues, while the later performance seen here might depend upon a late-stage convergence of the CD and IOVD signals involved in MID processing.

**Figure 7.**
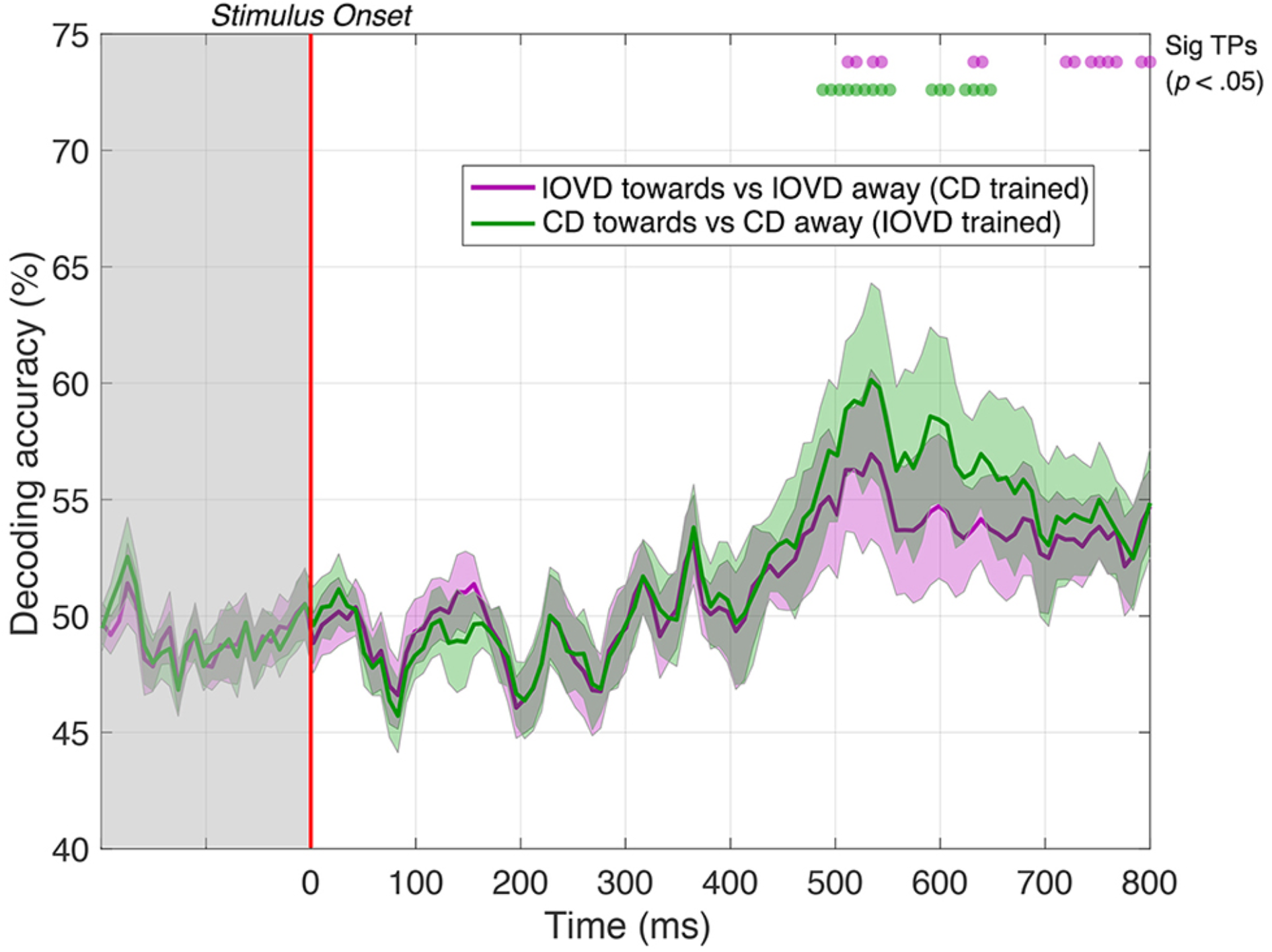
Pairwise decoding accuracy of a cross-trained decoder across 125 time points, comparing between towards and away directions for CD and IOVD stimuli. The purple line represents decoding accuracy for IOVD towards vs IOVD away after decoder is trained using CD direction data. The green line represents decoding accuracy for CD towards vs CD away after the decoder is trained using IOVD direction data. Purple and green dots indicate time points when the Bonferroni-corrected t-tests were significant for the 2 conditions (*p* < .05). Shaded error bars represent ±1SE of the bootstrapped mean (1000 iterations).

## Discussion

We have examined whether, and when, the pattern of EEG signals generated in response to CD- and IOVD-isolating stimuli moving towards or away in depth can be decoded using a multivariate pattern classifier. We find that both the 3D-motion direction (towards vs away) and MID cue type (CD vs IOVD) can be decoded based on the distinct pattern of neural responses measured across the scalp. We also show that CD motion direction decoding performance cannot be accounted for by the presence of static disparity information. Finally, and importantly, our data show that a cross-trained decoder (trained on one cue type, and then tested on the other) can decode EEG signals at relatively late stages of the EEG timecourse, suggesting a late-stage convergence in the processing of CD and IOVD information.

### Decoding MID direction

The decoder could distinguish between signals generated from stimuli moving towards or away in depth for both CD and IOVD stimuli. It has been well described that neurons in V1 and MT+ are selective for 2D-motion direction (Born & Bradley, 2005; Hubel & Wiesel, 1959, 1968) and disparity tuning (DeAngelis & Newsome, 1999; DeAngelis & Uka, 2003). Studies have also found evidence for MT+ neurons that are selective for 3D-motion direction, with both fMRI and psychophysical evidence for 3D direction-selective adaptation (Joo et al., 2016; Rokers et al., 2009, 2011). Notably, the peak for 3D-motion direction decoding occurred much later in the EEG timecourse than peak for MID-cue decoding. Brief stimuli produce a complex, temporally sustained pattern of cortical activity (including feed-forward and feedback signals, and the phase-resetting of endogenous rhythms) that persists for hundreds of milliseconds after stimulus offset (Bertamini et al., 2019; Mullinger et al., 2013, 2017; Stevenson et al., 2011). Further, 3D-motion direction signals take time to accumulate; motion direction can only be determined by integrating motion over time and the relatively late peak for 3D-motion direction decoding performance when compared to cue type discrimination is consistent with this (Baker & Bair, 2016). The relatively long time-scale required to decode 3D-motion direction after stimulus onset (from approximately 300ms onwards) is in line with what one might expect based on recent work of a similar vein (Bae & Luck, 2019), given the relatively weak percept of our stimulus, and the fact that the cortex must integrate its input for some time to make a complex decision about 3D-motion direction (Roitman & Shadlen, 2002). To note, we have decoded MID direction from responses recorded across the entire scalp, and the spatial resolution of EEG does not allow for localisation to specific visual areas. Thus, we do not suggest that our decoding MID direction is due to signals in MT, or V1 alone, although such visual regions might make some contribution to decoding performance. Importantly, our supplemental analysis showed that post-stimulus decoding could not be accounted for by different keyboard motor responses, as the decoder was unable to differentiate EEG signals between towards and away keyboard responses.

At the retinal level, the stimuli for the two motion directions (i.e., towards or away for a single cue type) are essentially identical at any given moment, and this might explain why motion direction decoding accuracy was relatively low when compared to CD vs IOVD cue decoding. Decoding 3D-motion direction in the case of the CD stimulus must depend on a mechanism that can compute a temporal derivative from disparity selective neurons. 3D-motion direction decoding in the case of the IOVD stimulus must be driven, at a minimum, by differential responses from monocular motion-selective neurons or by neurons that receive input from those populations. Most models of MID processing would posit that these 3D-motion direction selective neural populations are intermingled within visual areas involved in motion processing and so the ability to resolve these responses using EEG is intriguing (Czuba et al., 2014). A common suggestion within the field of EEG multivariate decoding is that neurons with different stimulus sensitivities are not co-mingled at random, but rather, are arranged in macroscale structures (such as columns or orientation domains), and their electrical signals might be differentiated – perhaps because of additional selectivity imposed by the curvature of the cortical surface (Cichy & Pantazis, 2017; da Silva, 2013; Hubel & Wiesel, 1974; Shoham et al., 1997; Sun et al., 2007). An alternative model might suggest that towards and away motion directions drive similar neural populations, however they might do so with differences in the timing or coherence. Overall, these data provide support for the existence of clusters of 3D-motion direction selective neurons, although we cannot identify the spatial scale of this clustering, or their broader cortical locus. It may be that our results are driven by 3D-motion direction selective neurons that are organised into columnar-scale structures within individual visual areas (much as 2D-direction selective neurons are), or that common populations of neurons process 3D-motion direction information but give rise to unique signals that differ in their timing, synchrony, or coherence.

### Decoding static disparity at early stages of the EEG time course

To examine the contribution of the time-averaged disparity information to CD direction decoding performance, we ran a second ‘static disparity’ experiment that used depth-fixed random dot stereogram stimuli with an identical distribution of starting positions, durations, and dot update rates to our original experiment. Although it was possible to decode static disparity information from the EEG signal at a few time points around 300 ms, decoding performance differed notably from CD direction decoding performance, which also rose above the statistical threshold around 300 ms, but then remained above chance for the remainder of the EEG timecourse. Static disparity information may drive the initial phases of CD decoding (i.e., the brief time point where decoding was possible for both experiments, around 300 ms), while later stages of MID decoding are attributable to motion through depth, rather than depth *per se.* The results of this static disparity experiment suggest that MID decoding is not based solely on responses from populations of motion-insensitive, disparity-selective neurons. However, this control cannot rule this possibility out entirely. Although human psychophysical observers are very accurate in discriminating between static stimuli with the maximum near and far disparities, and the discrimination performance for pairs of stimuli with zero disparity is necessarily at chance, in principle, the EEG signal classification may be qualitatively different for intermediate disparities that we have not tested. In future experiments, it would be of interest to compare psychophysical and EEG decoding performance in detail over this range for static stimuli.

### Decoding CD and IOVD cues

The decoder could accurately discriminate between CD and IOVD cues (pooled over 3D-motion direction) soon after stimulus onset. Here, decoding performance was high, with accuracy peaking at 84%. Similarly, we could decode CD and IOVD cues that were moving in the same direction in depth, indicating that most of the pooled decoding performance was driven by differences in the properties of CD and IOVD cues, rather than motion direction. Thus, our data show that the pattern of CD and IOVD signals are independent from each other at the level of the cortex, and that the intrinsic differences in the cue properties that give rise to the unique CD and IOVD retinal computations can be resolved downstream from the unique EEG signals that these cues provoke. Although we matched the low-level features of our CD and IOVD stimuli as closely as possible, they were, necessarily, different. CD and IOVD cues are thought to drive independent computations in the early visual system, and engage pathways with different spatial and temporal resolutions (Czuba et al., 2010; Shioiri et al., 2008; Wardle & Alais, 2013). A MID system that isolates CD and IOVD pathways will be necessarily reliant upon intrinsically different low-level stimulus properties. For example, CD monocular dot fields contain no monocular coherent motion energy and have a short lifetime, refreshing on every frame (i.e., every 16.6 ms across the two eyes), whereas IOVD dots travel short distances and have a relatively longer lifetime of 50 ms. Further, it may be that differences in the cortical signals generated by different eye-movement patterns to CD and IOVD cues contribute to between-cue decoding performance. Our data show that CD and IOVD cues give rise to distinct patterns of cortical signals, and the independence of these signals indicates that they would be processed via independent pathways. From this, one might hypothesise that neural computations involved in CD and IOVD processes are also kept independent from each other (Harris et al., 2008; Rokers et al., 2009).

### Cross trained decoding at late stages of the EEG time course

We cross-trained the decoder on the two different MID cues to examine the crucial question of whether decoding performance relies on shared or independent signals. We found that after being trained on CD towards and CD away signals, the decoder could accurately differentiate between IOVD towards and IOVD away signals, and vice-versa, at late stages of the EEG timecourse, beyond ~500 ms. This important result suggests that although the decoder must rely on unique CD and IOVD signals at the early and mid-stages of the EEG timecourse, there appears to be *shared information* at later stages of the timecourse. One possibility is that the cross-trained decoders performance at later stages depends on the convergence of signals involved in MID information processing. Our decoding performance was obtained using electrodes from across the full scalp and we do not attempt to spatially localise *where* this convergence occurs. One possibility is that a mechanism agnostic to differences in CD and IOVD cue properties, or possibly fuses both velocity and disparity signals together (Movshon & Newsome, 1996; Ponce et al., 2008). One candidate cortical location for this general stereomotion processing mechanism is the region in or around the human MT+ complex which includes cells that are known to be sensitive to both lateral motion defined from a variety of cues, as well as 3D-motion (Czuba et al., 2014; Héjja-Brichard et al., 2020; Huk, 2012; Kaestner et al., 2019; Likova & Tyler, 2007; Rokers et al., 2009; Sanada & DeAngelis, 2014). The temporal integration of 3D-motion signals has been shown to occur across hundreds of milliseconds, from ~150 to 1000 ms post-stimulus, with sensitivity to 3D-motion *increasing* across this integration period (Katz et al., 2015) and the behavioural integration of 3Dmotion signals across time is known to be relatively slow (when compared to lateral motion) (Brooks & Stone, 2006; Brooks & Stone, 2004; Harris et al., 2008; Harris & Watamaniuk, 1995, 1996; Huk, 2012; Norcia & Tyler, 1984; Richards, 1972). From this, combined with our relatively weak stimulus, and typical EEG dynamics, it is reasonable to assume that the convergence of MID signals can be measured ~500 ms post stimulus onset, although we reiterate, that the locus of such convergence cannot be identified in the current study.

## Conclusion

3D-motion direction (towards and away) and MID cues (CD and IOVD) can be decoded from the distinct pattern of EEG signals that they generate across the scalp. These data are consistent with reports suggesting the existence of clusters of 3D-motion direction selective neurons in visual cortex that may be organised into macroscale structures (Czuba et al., 2011; Joo et al., 2016). Further, we have shown that static disparity information can be decoded relatively early in the EEG timecourse, but it can play only a small, early role in CD direction decoding. Next, a MID system that isolates CD and IOVD necessarily relies on different low-level aspects stimulus properties, and we have shown that CD and IOVD cues can be resolved from the cortical signals that they provoke, suggesting that they are processed via two direction-specific independent pathways. Finally, results from a cross-trained decoder indicate that early stages of the EEG signal rely upon MID cue properties, while later stages of the signal rely on shared MID information. Overall, these data are the first to show that CD and IOVD cues that move towards and away in depth can be resolved and decoded using EEG, and that different aspects of MID-cues contribute to decoding performance along the EEG timecourse.

## Acknowledgements

The authors are grateful to Miaomiao Yu for assistance in setting up the EEG system. M.M.H and F.G.S share first-authorship.

## Supplementary Materials

### Behavioural responses

A paired samples t-test was used to examine differences in the proportion of correct responses when comparing between CD and IOVD stimuli (pooled across direction). As illustrated in **Figure S1A**, participants were significantly better at discriminating the direction of CD stimuli when compared to IOVD stimuli, *t*(19) = 2.269, *p* < .05, with participants 15.9% more accurate in identifying the direction of CD stimuli. Next, we split our data into the four stimulus conditions and a four-way repeated measures ANOVA was performed to assess whether there were differences in the proportion of correct responses between the four conditions. Mauchly’s test of sphericity was met (χ^2^(5) = 9.107, *p*= .107). The ANOVA found that there was a main effect of stimulus condition on the proportion of correct responses *F*(3,27) = 3.341, *p* < .05, η_p_^2^= .271, power = .693. As presented in **Figure S1B,** pairwise comparisons revealed that participants were more accurate at identifying CD towards responses when compared to IOVD towards responses (*p* < .05). No other comparisons were significant, suggesting that the increase in accuracy found in the pooled CD data when tested against the pooled IOVD data was primary driven by increased accuracy in the CD towards responses when compared to the IOVD conditions.

**Figure S1.**
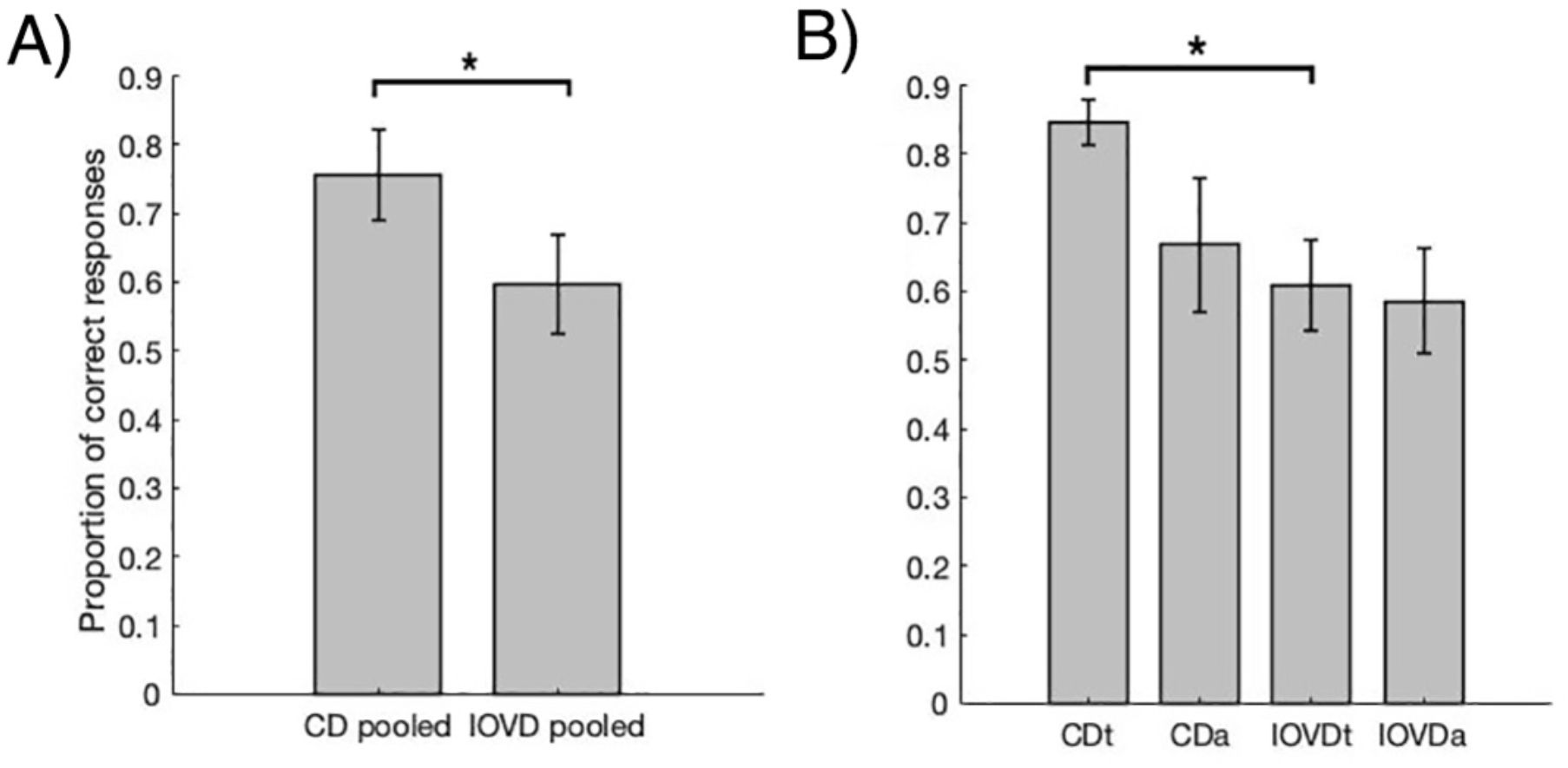
Bar plots of performance accuracy for identifying towards and away motion direction for MID stimuli. In panel A) we show the mean proportion of correct responses after data are pooled into CD and IOVD conditions and in B) we present the mean proportion of correct responses comparing across all four stimulus conditions. Error bars are ±1 SEM.

### Decoding keyboard responses: motor responses do not contribute to decoding performance

On each trial, participants were required to respond, via the keypad, whether they perceived the stimulus as moving towards or away in depth. The signals generated from pressing one of the two keys could, potentially, influence decoding accuracy. To address this, we ran the decoder on simulated data that differed only according to how participants responded on the keyboard during the trial. Here, we pooled our data into epochs labelled with either a ‘towards’ or an ‘away’ keyboard press response. This resulted in eight pools of data, for each participant: four in which the keyboard direction response matched the stimulus, four in which it was erroneous. We then performed a decoding analysis on keyboard button press (i.e. can we decode between when participant select towards or away on the keyboard?) using datasets that contained equal numbers of ‘towards’ and ‘away’ stimuli, drawn from these eight pools. Nine participants were included in this analysis. One participant was not included as they had only a single epoch in one of these pools (i.e. they were near perfect in identifying stimulus direction). For each of the eight data pools, we bootstrapped 210 epochs by taking four random epochs from each data pool and averaging them together. Thus, each of the eight data pools was now equal in size (210 simulated epochs), with four pools containing data made from ‘towards keyboard press’ epochs and the other four pools containing data made from ‘away keyboard press’ epochs. The subsequent analysis was identical to our main decoding analysis, repeatedly bootstrapping 21 ‘towards keypress’ epochs and 21 ‘away keypress’ epochs for each of 1000 iterations of the classifier.

The results (**Figure S2**) show that it is not possible to decode keyboard responses independently of the stimulus, with accuracy hovering around the 50% (chance) baseline across the 125 time points. These results suggest that stimulus decoding performance is likely to be driven by MID cue properties, rather than differences in signal from the motor cortex in response to keyboard button press.

**Figure S2.**
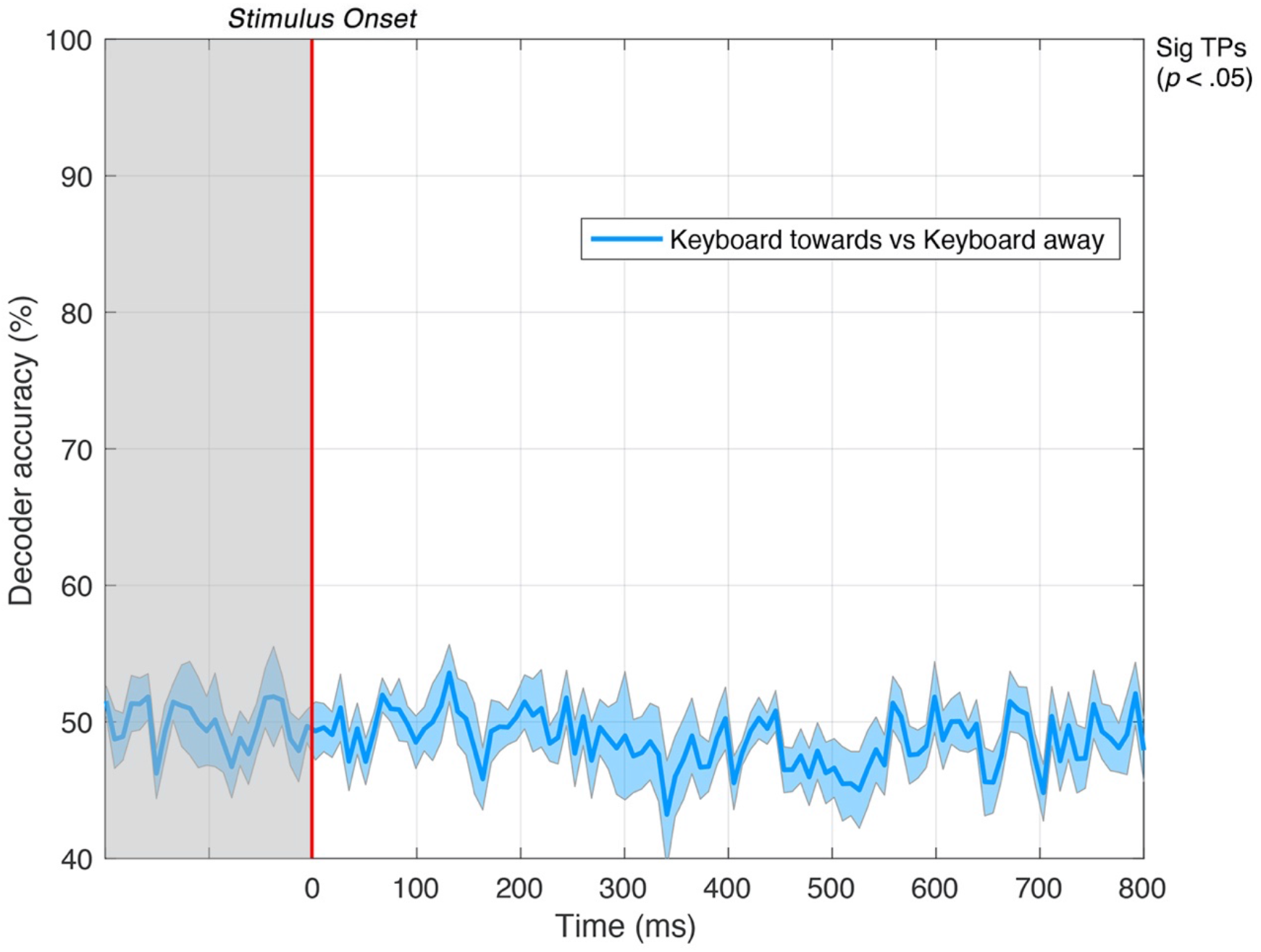
Pairwise decoding accuracy across 125 time points for keyboard response. Keyboard response (either pressing 2 on the keypad to indicate towards or 8 on the keypad to indicate away) cannot be accurately decoded: accuracy is at chance across all time points. Shaded error bars represent ±1SE of the bootstrapped mean (1000 iterations).

Thus, any post-stimulus decoding seen in this study cannot be accounted for by different keyboard motor responses as the decoder was unable to differentiate between towards and away keyboard responses. This has further implications. By decoding keyboard responses, we are effectively decoding the *percept* of 3D motion direction, that is, observers perceiving the stimulus either moving towards or away in depth (as well as occasional ‘button errors’). Thus, these results suggest that it is not possible to decode signals relating to the conscious percept of 3D motion direction for MID cues – although some brain regions (including motor cortex) do presumably carry such signals.

### Decoding the directional perception of MID based on behavioural responses

In an additional analysis we decoded the *conscious perception* of MID direction, that is, whether the classifier could distinguish between the pattern of EEG response based on whether a participant *perceived* the stimulus as moving either towards or away in depth (based on their keyboard response during the task), regardless of the actual stimulus direction. This differed from our ‘keyboard response’ analysis where each mean epoch was formed by drawing from each of the four stimulus conditions to average out any stimulus information, rather than drawing from CD or IOVD data pools alone.

Here, we partitioned our data based on MID stimulus type and keyboard response. We bootstrapped four epochs that were *perceived* as heading towards in depth and four epochs that were *perceived* as heading away in depth. This was completed for CD and IOVD cues separately. These eight trials were then averaged together to create a mean epoch (i.e. a ‘mean CD towards epoch’ is formed by averaging four epochs from CD towards data that were *correctly identified* as moving towards in depth and four epochs from CD away data that were *incorrectly identified* as the participant perceived them as moving towards in depth). This was to ensure that each mean epoch was formed of an equal count of correct and incorrectly perceived responses. Identical to our main analysis, this was repeated 21 times to create 21 mean epochs for each of the four conditions (CD perceived towards, CD perceived away, IOVD perceived towards, and IOVD perceived away) for each of the 1000 bootstrapped iterations of the decoder.

We found that decoding occurred around 50% chance (**Figure S3**). This suggests that successful decoding is based on the direction of the stimulus itself rather than its perceived direction or, critically, the preparation of the response-locked button press.

**Figure S3.**
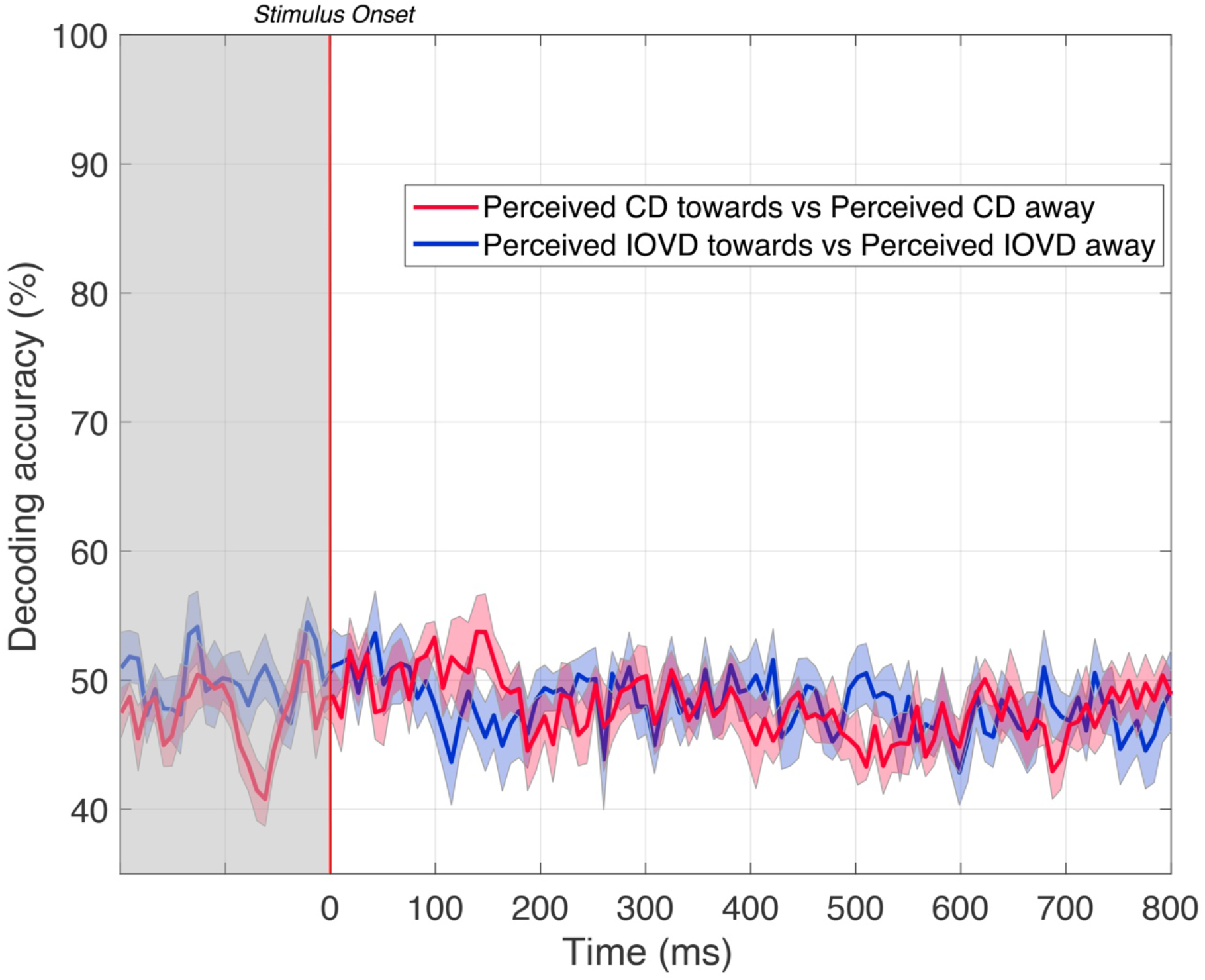
Pairwise decoding accuracies for comparing between data that was *perceived* as either moving towards or away in depth for CD and IOVD cues. Data is partitioned based on keyboard response, (*p* < .05). Shaded error bars represent ±1SE of the bootstrapped mean (1000 iterations).

### Scalp distributions

We decoded EEG responses using all 64 electrodes across the scalp. Here, we present the grand average common average referenced EEG response amplitudes for each stimulus condition, averaged across all participants. In **Figure S4** we present mean scalp maps of the EEG responses both CD and IOVD cues, averaged across three time periods. The scalp maps show a strong positive voltage towards the occipital cortex during early stages of the EEG time course. During middle stages, this changes to strong negative voltage near the occipital cortex, and a spreading of response across the scalp. This continues in later stages of EEG response, where the distribution is generally similar, but responses are weaker.

**Figure S4.**
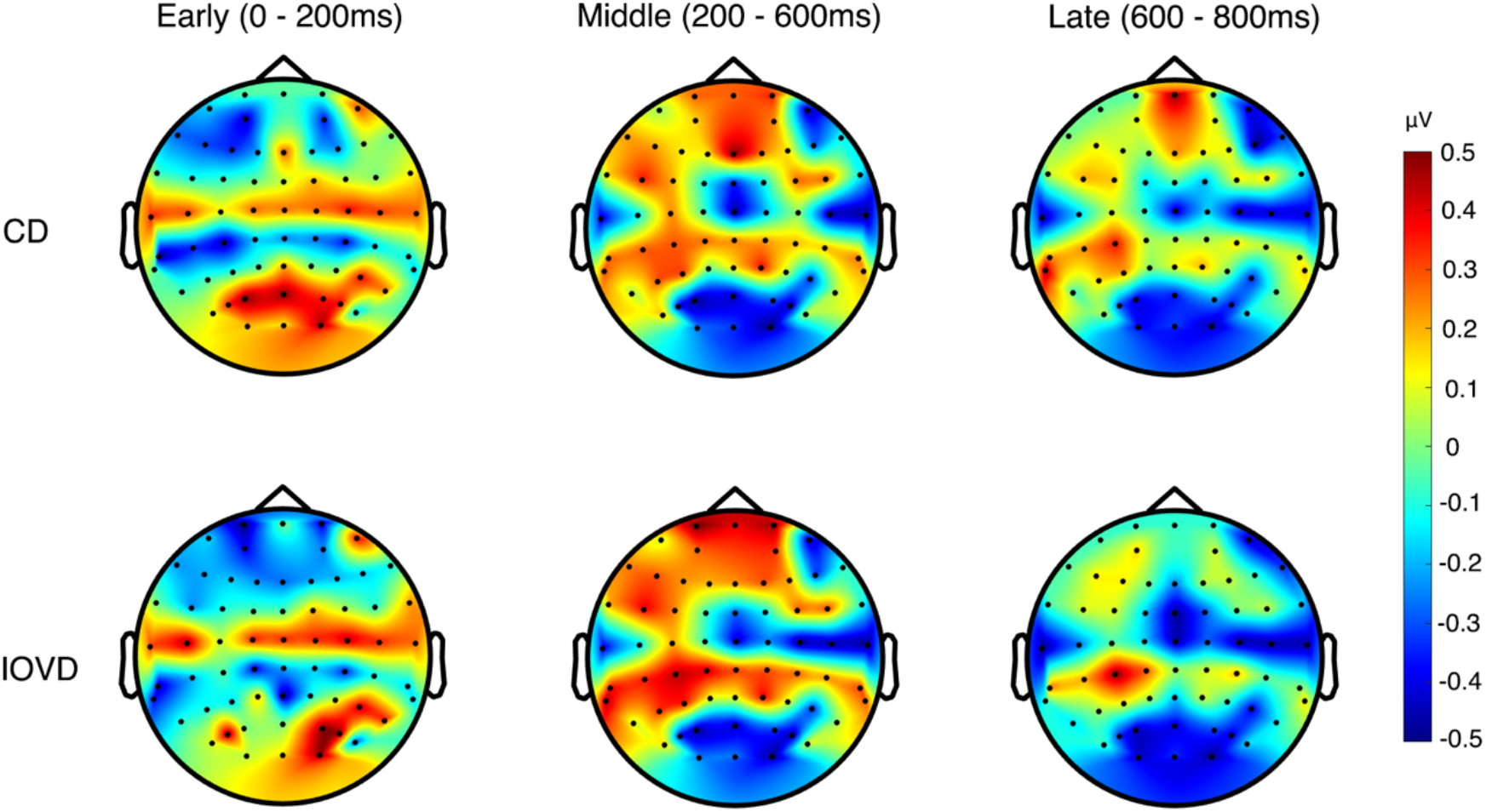
Scalp maps of mean common average referenced CD and IOVD (pooled across towards and away) EEG amplitudes are shown at three time periods – early (0 – 200ms post stimulus onset), middle (200-600ms post stimulus onset), and late (600 – 800ms post stimulus onset).

Similarly, in **Figure S5** we present scalp maps of the mean EEG responses both towards and away cues, averaged across three time periods. Again, these scalp maps show an initial strong positive voltage towards near the occipital cortex during early stages of the EEG time course, in middle stages this changes to a strong negative voltage near the occipital cortex, and the response distribution spreads across the scalp. This continues in later stages of EEG response, albeit with weaker responses.

**Figure S5.**
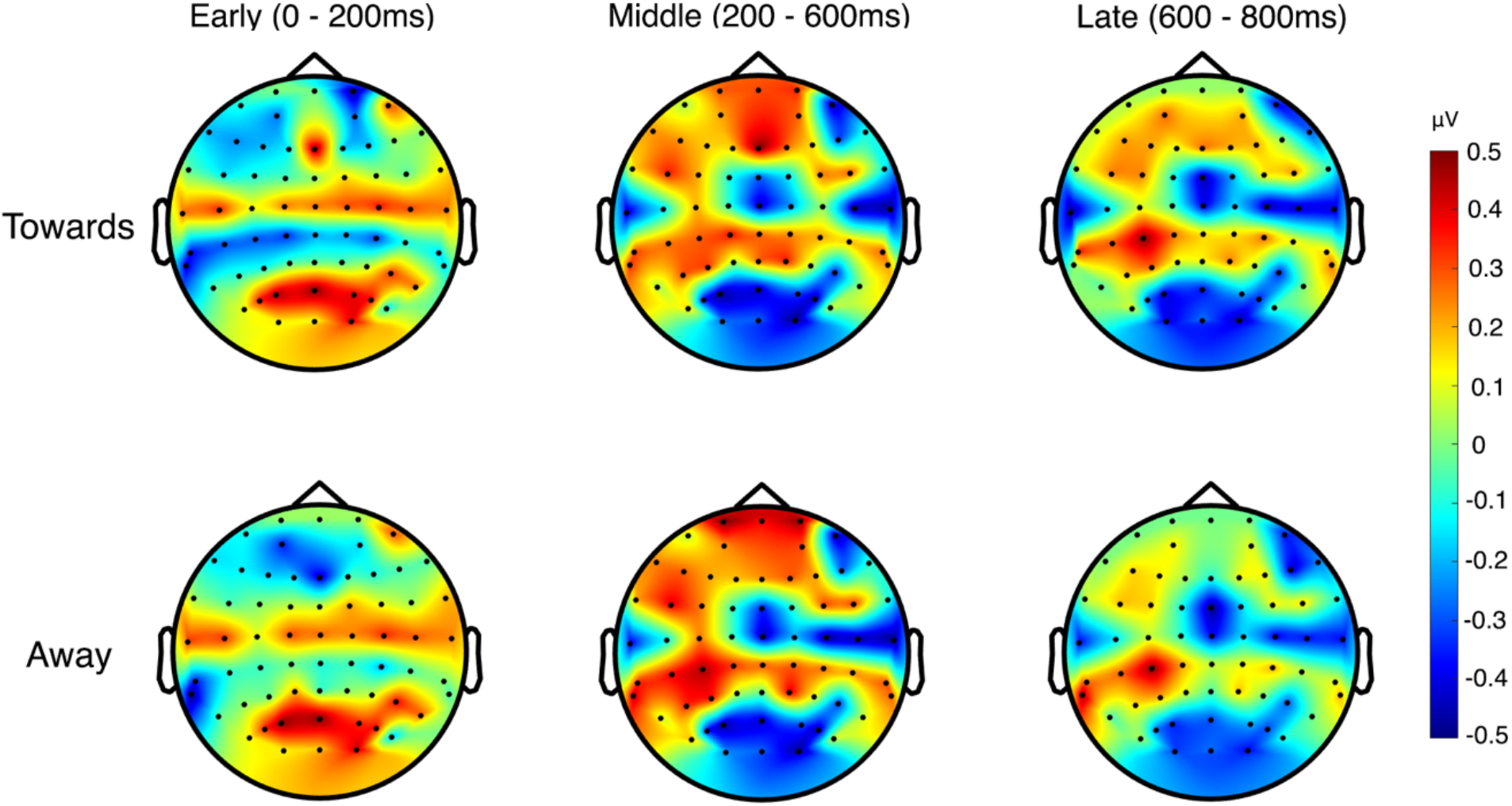
Scalp montages of mean common average referenced towards and away (pooled across CD and IOVD) EEG amplitudes are shown at three time periods – early (0 – 200ms post stimulus onset), middle (200-600ms post stimulus onset), and late (600 – 800ms post stimulus onset).

### Decoding validation from shuffled labels

**Figure S6.**
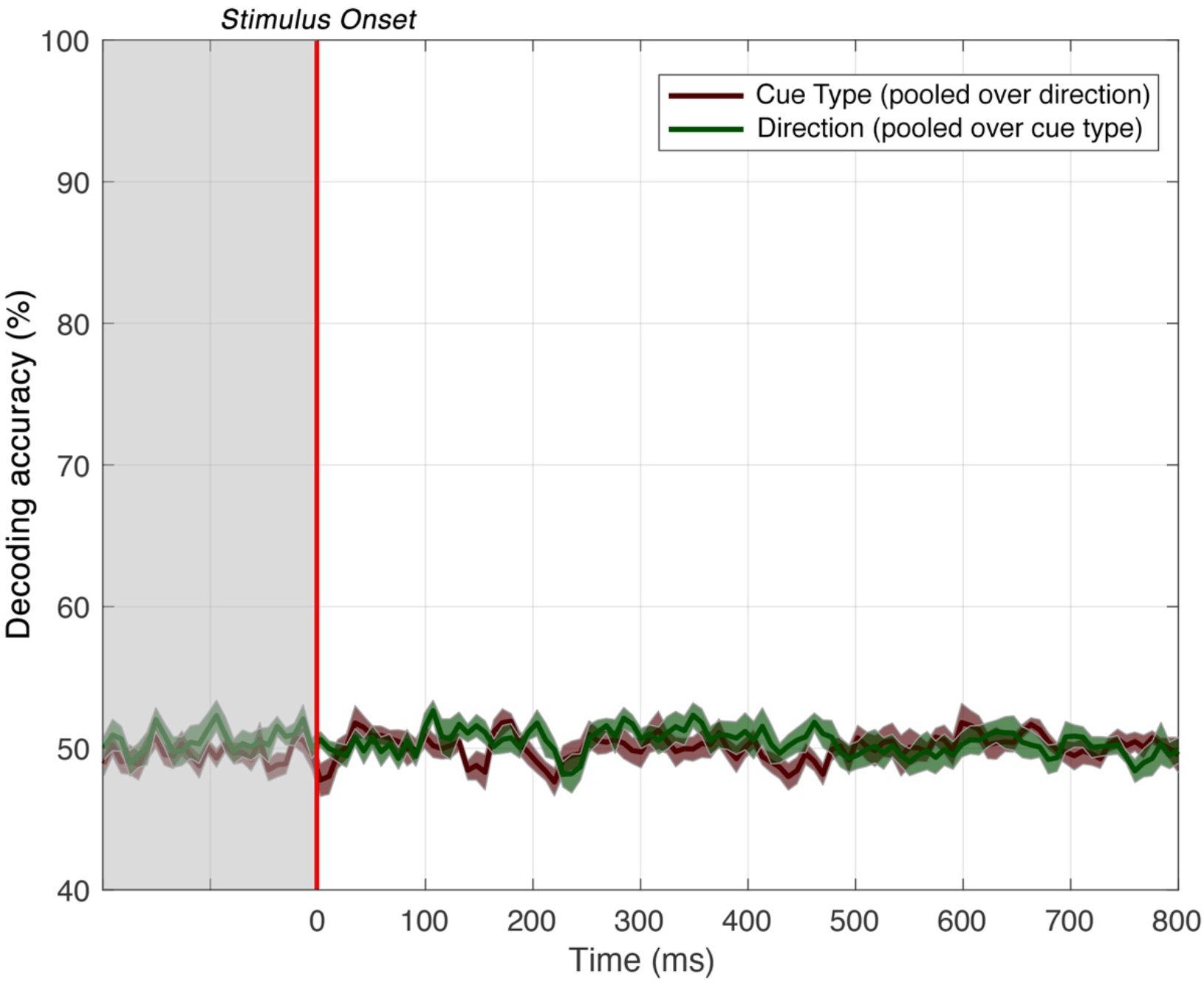
Shuffled label pairwise decoding accuracies for cue type (pooled over direction) in red and direction (pooled over cue type) in green. After shuffling labels, the decoding accuracy of both comparisons fell around the 50% (chance) baseline, confirming that the SVM was working as expected (*p* < .05). Shaded error bars represent ±1SE of the bootstrapped mean (1000 iterations).

**Figure S7.**
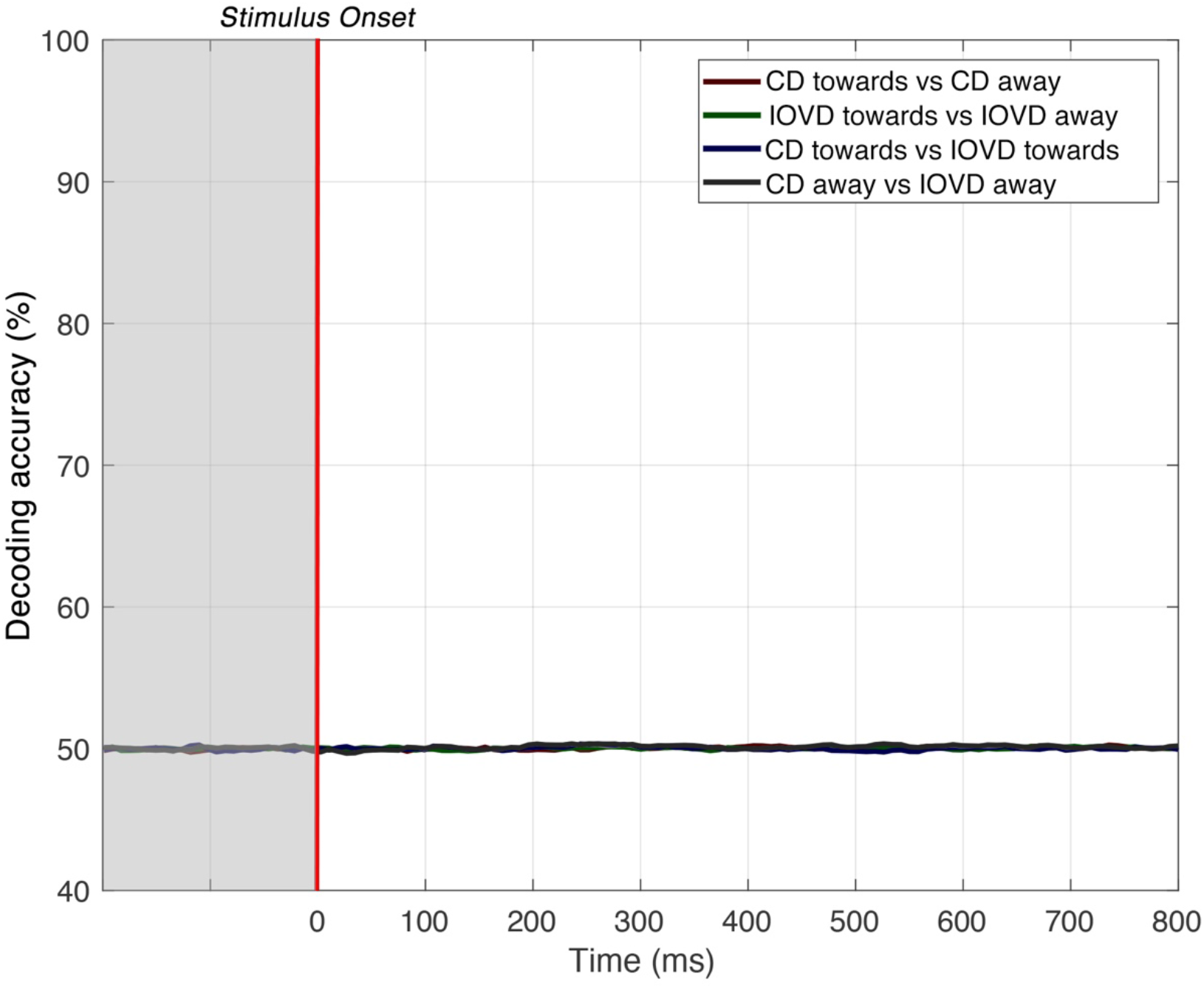
Shuffled label pairwise decoding accuracies for within cue and within direction conditions. After shuffling labels, accuracy for all four conditions was virtually identical, falling around the 50% (chance) baseline, confirming that the SVM was working as expected (p < .05). Shaded error bars represent ±1SE of the bootstrapped mean (1000 iterations).

### Extended results of CD and IOVD discrimination within motion direction

**Figure S8.**
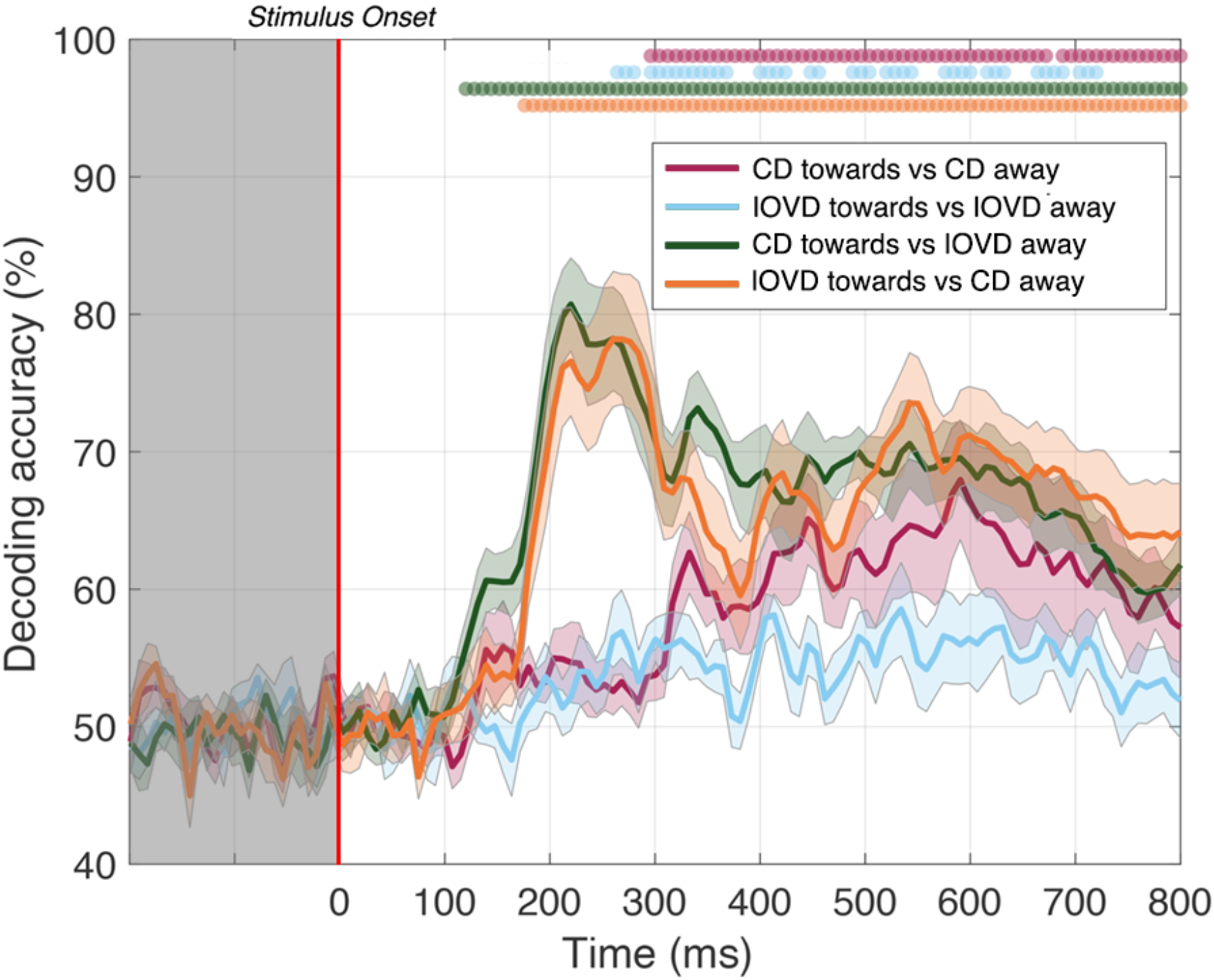
Pairwise decoding accuracy across 125 time points for four stimulus conditions. We included CD towards vs IOVD away and IOVD towards vs CD away to compare with decoding accuracy in **Figure 5**. These new comparisons do not particularly differ from those in **Figure 5**, suggesting that direction decoding cannot be disentangled from cue decoding, and that motion direction discrimination might be a shared process between cue types. Red, light blue, green, and orange dots indicate time points when the Bonferroni-corrected t-tests were significant for each condition (p < .05) and the coloured ticks on the x-axis represent the time point of peak decoding performance for each stimulus condition. Shaded error bars represent ±1SE of the bootstrapped mean (1000 iterations).

## References

Bae, G.-Y., & Luck, S. J.(2019). Decoding motion direction using the topography of sustained ERPs and alpha oscillations. NeuroImage, 184, 242–255. https://doi.org/10.1016/j.neuroimage.2018.09.029

Baker, D. H., Kaestner, M., & Gouws, A. D. (2016). Measurement of crosstalk in stereoscopic display systems used for vision research. J Vis, 16(15), 14. https://doi.org/10.1167/16.15.14

Baker, P. M., & Bair, W. (2016). A model of binocular motion integration in MT neurons. Journal of Neuroscience, 36(24), 6563–6582. https://doi.org/10.1523/JNEUROSCI.3213-15.2016

Bertamini, M., Rampone, G., Oulton, J., Tatlidil, S., & Makin, A. D. J. (2019). Sustained response to symmetry in extrastriate areas after stimulus offset: An EEG study. Scientific Reports, 9(1), 4401. https://doi.org/10.1038/s41598-019-40580-z

Born, R. T., & Bradley, D. C. (2005). Structure and function of visual area MT. Annual Review of Neuroscience, 28, 157–189. https://doi.org/10.1146/annurev.neuro.26.041002.131052

Boser, B., Guyon, I., & Vapnik, V. (1992). A Training Algorithm for Optimal Margin Classifiers. Proceedings of the 5th Annual ACM Workshop on Computational Learning Theory, 144–152. https://doi.org/10.1145/130385.130401

Brainard, D. H. (1997). The Psychophysics Toolbox. Spatial Vision, 10(4), 433–436. https://doi.org/10.1163/156856897X00357

Brooks, K. R., & Stone, L. S. (2006). Stereomotion suppression and the perception of speed: Accuracy and precision as a function of 3D trajectory. Journal of Vision, 6(11), 6–6. https://doi.org/10.1167/6.11.6

Brooks, Kevin R, & Stone, L. S. (2004). Stereomotion speed perception: Contributions from both changing disparity and interocular velocity difference over a range of relative disparities. Journal of Vision, 4(12), 1061–1079. https://doi.org/10:1167/4.12.6

Bullier, J. (2001). Integrated model of visual processing. Brain Research Reviews, 36(2–3), 96–107. https://doi.org/10.1016/S0165-0173(01)00085-6

Casagrande, V. A., & Boyd, J. D. (1996). The neural architecture of binocular vision. Eye, 10(2), 153–160. https://doi.org/10.1038/eye.1996.40

Chang, C., & Lin, C. (2011). LIBSVM -- A Library for Support Vector Machines. ACM Transactions on Intelligent Systems and Technology, 2(3), 1–27. https://doi.org/10.1145/1961189.1961199

Cichy, R. M., & Pantazis, D. (2017). Multivariate pattern analysis of MEG and EEG: A comparison of representational structure in time and space. NeuroImage, 158, 441–454.

Cortes, C., & Vapnik, V. (1995). Support-Vector Networks. Machine Learning, 20, 273–297. https://doi.org/10.1023/A:1022627411411

Cottereau, B. R., Ales, J. M., & Norcia, A. M. (2014). The evolution of a disparity decision in human visual cortex. NeuroImage, 92, 193–206. https://doi.org/10.1016/j.neuroimage.2014.01.055

Cottereau, B. R., McKee, S. P., Ales, J. M., & Norcia, A. M. (2011). Disparity-tuned population responses from human visual cortex. Journal of Neuroscience, 31(3), 954–965.

Cottereau, B. R., McKee, S. P., & Norcia, A. M. (2013). Dynamics and cortical distribution of neural responses to 2D and 3D motion in human. Journal of Neurophysiology. https://doi.org/10.1152/jn.00549.2013

Cumming, B. G., & Parker, A. J. (1994). Binocular mechanisms for detecting motion-in-depth. Vision Research, 34(4), 483–495.

Czuba, T. B., Huk, A. C., Cormack, L. K., & Kohn, A. (2014). Area MT encodes three-dimensional motion. J Neurosci, 34(47), 15522–15533. https://doi.org/10.1523/JNEUROSCI.1081-14.2014

Czuba, T. B., Rokers, B., Guillet, K., Huk, A. C., & Cormack, L. K. (2011). Three-dimensional motion aftereffects reveal distinct direction-selective mechanisms for binocular processing of motion through depth. J Vis, 11(10), 18. https://doi.org/10.1167/11.10.18

Czuba, T. B., Rokers, B., Huk, A. C., & Cormack, L. K. (2010). Speed and eccentricity tuning reveal a central role for the velocity-based cue to 3D visual motion. J Neurophysiol, 104(5), 2886–2899. https://doi.org/10.1152/jn.00585.2009

da Silva, F. L. (2013). EEG and MEG: Relevance to Neuroscience. Neuron, 80(5), 1112–1128.

DeAngelis, G. C., & Newsome, W. T. (1999). Organization of disparity-selective neurons in macaque area MT. The Journal of Neuroscience, 19(4), 1398–1415.

DeAngelis, G. C., & Uka, T. (2003). Coding of Horizontal Disparity and Velocity by MT Neurons in the Alert Macaque. Journal of Neurophysiology, 89(2), 1094–1111. https://doi.org/10.1152/jn.00717.2002

Grootswagers, T., Wardle, S. G., & Carlson, T. A. (2017). Decoding Dynamic Brain Patterns from Evoked Responses: A Tutorial on Multivariate Pattern Analysis Applied to Time Series Neuroimaging Data. Journal of Cognitive Neuroscience, 29(4), 677–697. https://doi.org/10.1162/jocn_a_01068

Harris, J. M., Nefs, H. T., & Grafton, C. E. (2008). Binocular vision and motion-in-depth. Spatial Vision, 21(6), 531–547.

Harris, J. M., & Watamaniuk, S. N. (1995). Speed discrimination of motion-in-depth using binocular cues. Vision Research, 35(7), 885–896.

Harris, J. M., & Watamaniuk, S. N. (1996). Poor speed discrimination suggests that there is no specialized speed mechanism for cyclopean motion. Vision Research, 36(14), 2149–2157.

Héjja-Brichard, Y., Rima, S., Rapha, E., Durand, J., & Cottereau, B. R. (2020). Stereomotion processing in the non-human primate brain. Cerebral Cortex. https://doi.org/10.1101/638155

Howard, I. P., & Rogers, B. J. (2002). Seeing in depth (Vol. 2). I. Porteous.

Hubel, D. H., & Wiesel, T. N. (1959). Receptive fields of single neurones in the cat’s striate cortex. Journal of Physiology, 148(3), 574–591. https://doi.org/10.1113/jphysiol.1959.sp006308

Hubel, D. H., & Wiesel, T. N. (1968). Receptive fields and the functional architecture of monkey striate cortex. Journal of Physiology, 195(1), 215–243. https://doi.org/10.1113/jphysiol.1968.sp008455

Hubel, D. H., & Wiesel, T. N. (1974). Sequence regularity and geometry of orientation columns in the monkey striate cortex. The Journal of Comparative Neurology, 158, 267–293.

Huk, A. C. (2012). Multiplexing in the primate motion pathway. Vision Research, 62, 173–180.

Joo, S. J., Czuba, T. B., Cormack, L. K., & Huk, A. C. (2016). Separate perceptual and neural processing of velocity-and disparity-based 3D motion signals. J Neurosci, 36(42), 10791–10802. https://doi.org/10.1523/JNEUROSCI.1298-16.2016

Kaestner, M., Maloney, R. T., Wailes-Newson, K. H., Bloj, M., Harris, J. M., Morland, A. B., & Wade, A. R. (2019). Asymmetries between achromatic and chromatic extraction of 3D motion signals. Proceedings of the National Academy of Sciences, 116(27), 13631–13640. https://doi.org/10.1073/pnas.1817202116

Katz, L. N., Hennig, J. A., Cormack, L. K., & Huk, A. C. (2015). A Distinct Mechanism of Temporal Integration for Motion through Depth. Journal of Neuroscience, 35(28), 10212–10216. https://doi.org/10.1523/JNEUROSCI.0032-15.2015

Lamme, V. A. F., & Roelfsema, P. R. (2000). The distinct modes of vision offered by feedforward and recurrent processing. Trends in Neurosciences, 23(11), 571–579. https://doi.org/10.1016/S0166-2236(00)01657-X

Likova, L. T., & Tyler, C. W. (2007). Stereomotion processing in the human occipital cortex. NeuroImage, 38(2), 293–305. https://doi.org/10.1016/j.neuroimage.2007.06.039

Maloney, R. T., Kaestner, M., Bruce, A., Bloj, M., Harris, J. M., & Wade, A. R. (2018). Sensitivity to velocity-and disparity-based cues to motion-In-depth with and without spared stereopsis in binocular visual impairment. Investigative Ophthalmology & Visual Science, 59(11), 4375–4383. https://doi.org/10.1167/iovs.17-23692

Maris, E., & Oostenveld, R. (2007). Nonparametric statistical testing of EEG-and MEG-data. Journal of Neuroscience Methods, 164(1), 177–190. https://doi.org/10.1016/j.jneumeth.2007.03.024

Movshon, J. A., & Newsome, W. T. (1996). Visual response properties of striate cortical neurons projecting to area MT in macaque monkeys. Journal of Neuroscience, 16(23), 7733–7741.

Mullinger, K. J., Cherukara, M. T., Buxton, R. B., Francis, S. T., & Mayhew, S. D. (2017). Post-stimulus fMRI and EEG responses: Evidence for a neuronal origin hypothesised to be inhibitory. NeuroImage, 157, 388–399. https://doi.org/10.1016/j.neuroimage.2017.06.020

Mullinger, K. J., Mayhew, S. D., Bagshaw, A. P., Bowtell, R., & Francis, S. T. (2013). Poststimulus undershoots in cerebral blood flow and BOLD fMRI responses are modulated by poststimulus neuronal activity.Proceedings of the National Academy of Sciences, 110(33), 13636–13641. https://doi.org/10.1073/pnas.1221287110

Norcia, A. M., & Tyler, C. W. (1984). Temporal frequency limits for stereoscopic apparent motion processes. Vision Research, 24(5), 395–401.

Ponce, C. R., Lomber, S. G., & Born, R. T. (2008). Integrating motion and depth via parallel pathways. Nature Neuroscience, 11(2), 216–223. https://doi.org/10.1038/nn2039

Rashbass, C., & Westheimer, G. (1961). Independence of conjugate and disjunctive eye movements. J Physiol, 159, 361–364.

Regan, D. (1993). Binocular correlates of the direction of motion in depth. Vision Research, 33(16), 2359–2360. https://doi.org/10.1016/0042-6989(93)90114-C

Richards, W. (1972). Response Functions for Sine-and Square-Wave Modulations of Disparity. Journal of the Optical Society of America, 62(7), 907. https://doi.org/10.1364/JOSA.62.000907

Roitman, J. D., & Shadlen, M. N. (2002). Response of Neurons in the Lateral Intraparietal Area during a Combined Visual Discrimination Reaction Time Task. The Journal of Neuroscience, 22(21), 9475–9489. https://doi.org/10.1523/JNEUROSCI.22-21-09475.2002

Rokers, B., Cormack, L. K., & Huk, A. C. (2009). Disparity-and velocity-based signals for three-dimensional motion perception in human MT+. Nature Neuroscience, 12(8), 1050–1055.

Rokers, B., Czuba, T. B., Cormack, L. K., & Huk, A. C. (2011). Motion processing with two eyes in three dimensions. J Vis, 11(2). https://doi.org/10.1167/11.2.10

Sakano, Y., Allison, R. S., & Howard, I. P. (2012). Motion aftereffect in depth based on binocular information. Journal of Vision, 12(1), 11–11. https://doi.org/10.1167/12.1.11

Sanada, T. M., & DeAngelis, G. C. (2014). Neural representation of motion-in-depth in area MT. J Neurosci, 34(47), 15508–15521. https://doi.org/10.1523/JNEUROSCI.1072-14.2014

Shioiri, S., Nakajima, T., Kakehi, D., & Yaguchi, H. (2008). Differences in temporal frequency tuning between the two binocular mechanisms for seeing motion in depth. J Opt Soc Am A Opt Image Sci Vis, 25(7), 1574–1585.

Shioiri, S., Saisho, H., & Yaguchi, H. (2000). Motion in depth based on inter-ocular velocity differences. Vision Res, 40(19), 2565–2572.

Shoham, D., Hübener, M., Schulze, S., Grinvald, A., & Bonhoeffer, T. (1997). Spatiotemporal Frequency Domains and Their Relation to Cytochrome Oxidase Staining in Cat Visual Cortex. Nature, 385, 529–533.

Stevenson, C. M., Brookes, M. J., & Morris, P. G. (2011). β-Band correlates of the fMRI BOLD response. Human Brain Mapping, 32(2), 182–197. https://doi.org/10.1002/hbm.21016

Sun, P., Ueno, K., Waggoner, R. A., Gardner, J. L., Tanaka, K., & Cheng, K. (2007). A temporal frequency–dependent functional architecture in human V1 revealed by high-resolution fMRI. Nature Neuroscience, 10(11), 1404–1406. https://doi.org/10.1038/nn1983

Wang, L. (2018). Support Vector Machines: Theory and Applications (Studies in Fuzziness and Soft Computing). Springer-Verlag.

Wardle, S. G., & Alais, D. (2013). Evidence for speed sensitivity to motion in depth from binocular cues. Journal of Vision, 13(1). https://doi.org/10.1167/13.1.17

